# Biofunctional 2D Graphitic Carbon Nitride-Hydrogel Heterointerfaces for Electrochemical Detection of Interleukin-6 toward Septic Cardiomyopathy Diagnostics in Clinical Biofluids

**DOI:** 10.64898/2026.05.25.727622

**Authors:** Pranjal Agarwal, Amit K. Yadav, Ankur Singh, Sumit K. Yadav, N V S Praneeth, Dhiraj Bhatia

**Affiliations:** Department of Biological Sciences and Engineering, Indian Institute of Technology Gandhinagar (IITGN), Near Palaj, Gandhinagar, Gujarat 382355, India; Health Technology Cluster, School of Health Sciences and Technology, University of Petroleum and Energy Studies (UPES), Dehradun, Uttarakhand, India-248007; Department of Chemistry, Indian Institute of Technology Gandhinagar (IITGN), Near Palaj, Gandhinagar, Gujarat 382355, India

**Author notes:** **Corresponding Authors:** Dr. Amit K. Yadav; Email-, Prof. Dhiraj Bhatia. Authors Contributed Equally to this work (Pranjal Agarwal, Amit K. Yadav).

**Keywords:** Electrochemical Aptasensing, Graphitic Carbon Nitride, Chitosan Hydrogel, Septic cardiomyopathy, Interleukin-6

## Abstract

Interleukin-6 (IL-6) is a key pro-inflammatory cytokine closely associated with sepsis progression and septic cardiomyopathy (SCM), a severe clinical condition characterized by acute cardiac dysfunction and high mortality in critically ill patients. Rapid and sensitive monitoring of IL-6 in clinical biofluids is therefore crucial for early diagnosis, disease prognosis, and timely therapeutic intervention in emergency healthcare settings. Herein, we report a biofunctional, label-free electrochemical aptasensor based on a graphitic carbon nitride-incorporated chitosan hydrogel-modified gold screen-printed electrode (MCH/Apt-IL-6/g-C_3_N_4_@CS/Au-SPE) for ultrasensitive detection of IL-6 in clinical biofluids. The electroactive g-C_3_N_4_@CS hydrogel heterointerface was engineered via electrostatic interactions between the negatively charged surface functionalities of two-dimensional graphitic carbon nitride (g-C_3_N_4_) and the protonated amino groups (-NH_3_^+^) of chitosan (CS), yielding a porous, conductive, and biocompatible sensing matrix with enhanced aptamer immobilization and accelerated electron-transfer kinetics. Biocompatibility evaluation using MTT assay and confocal fluorescence imaging demonstrated that the hydrogel maintained excellent cellular compatibility at 5 mg/mL, preserving normal cytoskeletal organization, mitochondrial integrity, and nuclear morphology, while higher concentrations induced cellular stress responses. Under optimized experimental conditions, the developed aptasensor exhibited outstanding analytical performance with an ultrawide linear detection range from 1 fg/mL to 10 ng/mL, a high sensitivity of 2.162 μA/[log_10_(ng/mL)] cm^-2^, a low detection limit of 0.460 pg/mL, and excellent linearity (R^2^ = 0.979). In addition, the sensor demonstrated remarkable selectivity toward common biological interferents, including ascorbic acid, cysteine, glucose, glycine, and urea, together with excellent reproducibility (RSD = 1.139%). Validation studies performed in spiked human serum samples further confirmed the reliability and practical applicability of the proposed sensing platform for rapid clinical analysis. Owing to its label-free detection strategy, disposable electrode format, high sensitivity, and favorable biocompatibility, the developed g-C_3_N_4_-hydrogel heterointerface-based aptasensor represents a promising next-generation platform for early septic cardiomyopathy diagnostics, inflammatory biomarker monitoring, and point-of-care electrochemical biosensing applications.

## 1. INTRODUCTION

Sepsis is a life-threatening condition characterized by a dysregulated host response to infection, often leading to multiple organ dysfunction and high mortality rates^1–3^. Among its severe complications, septic cardiomyopathy (SCM) has emerged as a major contributor to adverse clinical outcomes, apparently as reversible cardiac dysfunction, reduced ejection fraction, and impaired ventricular contractility^4–7^. The complex and multifactorial pathophysiology of SCM involves inflammatory cascades, oxidative stress, mitochondrial dysfunction, and altered calcium homeostasis^5,8,9^. Despite advances in critical care, the early diagnosis of SCM remains challenging due to the lack of specific, rapid diagnostic tools, underscoring the urgent need for reliable biomarkers to facilitate timely intervention and improve patient prognosis^6,9,10^. Inflammatory mediators play a central role in the progression of sepsis and SCM, among which interleukin-6 (IL-6) has been widely recognized as a key pro-inflammatory cytokine^11–13^. IL-6 is rapidly produced in response to infection and tissue injury, and its elevated levels have been strongly correlated with disease severity, cardiac dysfunction, and mortality in septic patients^12,14,15^. Owing to its early release and clinical relevance, IL-6 serves as a valuable biomarker for both diagnosis and prognosis in sepsis-associated conditions^15,16^. However, the dynamic and often low-concentration expression of IL-6 in biological fluids necessitates highly sensitive, rapid, and reliable detection methods^17–19^. Conventional analytical techniques, although widely used, face significant limitations in meeting these requirements^20–22^. Traditional methods for IL-6 detection, such as enzyme-linked immunosorbent assays (ELISA), polymerase chain reaction (PCR), and other immunoassays, are often labor-intensive, time-consuming, and require sophisticated instrumentation and trained personnel^23^. These limitations limit their applicability for real-time monitoring and point-of-care (POC) diagnostics, which are crucial for managing acute conditions such as sepsis^24,25^. In recent years, there has been growing interest in developing portable, rapid diagnostic platforms that can provide timely, accurate biomarker detection^26,27^. In this context, biosensor-based technologies have emerged as promising alternatives, offering advantages such as high sensitivity and specificity, rapid response, and the potential for miniaturization^25^. Biosensors are analytical devices that integrate a biological recognition element with a physicochemical transducer to convert biological interactions into measurable signals^28–30^. The integration of nanotechnology into biosensor design has significantly enhanced their performance, leading to the development of next-generation nanobiosensors^31–33^. These systems exploit the unique physicochemical properties of nanomaterials, including high surface-to-volume ratio, enhanced electrical conductivity, and tunable surface functionalities, to achieve improved signal transduction and ultra-low detection limits^34,35^. Hydrogel-based biosensors, in particular, have attracted considerable attention due to their ability to combine the advantageous properties of multiple materials, thereby enhancing sensitivity, selectivity, and stability^36^. Such platforms have shown great potential for detecting clinically relevant biomarkers, especially in complex biological matrices^37^.

Among various nanomaterials, graphitic carbon nitride (g-C_3_N_4_) has emerged as a promising biosensing material due to its unique structural and electronic properties^38–40^. As a metal-free, two-dimensional polymeric material, g-C_3_N_4_ exhibits high chemical stability, excellent biocompatibility, and a moderate band gap that facilitates efficient electron transfer^41,42^. Its layered structure provides a large surface area for biomolecule immobilization, while its rich nitrogen content enables facile functionalization^43,44^. These features make g-C_3_N_4_ particularly attractive for enhancing the sensitivity and performance of electrochemical and optical biosensors. However, its practical application is often limited by issues such as poor dispersibility and aggregation, which can reduce the effective surface area and hinder its performance^45^. To overcome these limitations, the incorporation of biocompatible polymers such as chitosan has been widely explored. Chitosan (CS), a naturally derived polysaccharide obtained from chitin, is known for its excellent biocompatibility, biodegradability, and non-toxicity^46–48^. It possesses abundant amino and hydroxyl functional groups, which facilitate the immobilization of biomolecules and enhance the stability of biosensing platforms^49–51^. Additionally, chitosan exhibits excellent film-forming properties and can readily form hydrogels with a three-dimensional network, providing a supportive matrix for nanomaterials and biological components^49,52^. The combination of chitosan with nanomaterials not only improves dispersion and stability but also creates a favorable microenvironment for efficient signal transduction and biomolecular interactions^46,51,53^.

The integration of nanomaterials with hydrogel systems has opened new avenues in biosensor design. Hydrogels exhibit enhanced porosity, mechanical stability, and mass transport properties, enabling improved loading of biorecognition elements and efficient interaction with target analytes^54^. Recent advances demonstrate that such hybrid systems can achieve superior sensitivity and selectivity for biomarker detection, particularly in complex biological environments^55^. Furthermore, integrating nanobiosensors into advanced platforms, such as wearable devices and organ-on-chip systems, enables real-time monitoring of cytokines and other biomarkers under physiological conditions^56^.

In this regard, the development of g-C_3_N_4_@CS hydrogel-based aptasensor represents a promising strategy for the design of advanced biosensing platforms. The synergistic integration of g-C_3_N_4_ and chitosan combines the excellent electrical and catalytic properties of the former with the biocompatibility and structural versatility of the latter. The hydrogel matrix offers a high surface area, enhanced porosity, and efficient mass transport, enabling improved loading of biorecognition elements and rapid analyte diffusion. Such hybrid systems are particularly well-suited for detecting low-abundance biomarkers such as IL-6, where sensitivity and selectivity are critical. In this study, we report the development of a novel g-C_3_N_4_@CS hydrogel-based biosensor for sensitive and selective detection of IL-6, a biomarker of septic cardiomyopathy. The proposed platform leverages the synergistic properties of the hydrogel to achieve enhanced electrochemical performance, rapid response, and improved detection limits. By addressing the limitations of conventional diagnostic methods, this work aims to provide a reliable and efficient tool for early diagnosis and monitoring of sepsis-associated cardiac dysfunction. The findings of this study highlight the potential of hydrogel-based biosensors to advance point-of-care diagnostics and improve clinical outcomes in critical care settings.

## 2. MATERIALS AND METHODS

### 2.1 Reagents and chemicals

All chemical compounds and reagents employed in the current study, including ethanol (CH_3_CH_2_OH, ≥99.9%), isopropyl alcohol (IPA, ≥99.9%), potassium hexacyanoferrate(III) (K_3_[Fe(CN)_6_], 99%), potassium hexacyanoferrate(II) trihydrate (K_4_[Fe(CN)_6_]·3H_2_O, ≥99.95%), 6-mercapto-1-hexanol (MCH, ≥97%), tris(2-carboxyethyl)phosphine hydrochloride (TCEP, ≥98%), chitosan (medium molecular weight), sodium phosphate dibasic dihydrate (NaH_2_PO_4_.2H_2_O, ≥99.0%), sodium phosphate monobasic anhydrous (NaH_2_PO_4_, ≥98%), sodium chloride (NaCl, ≥99.0%), ascorbic acid (C_6_H_8_O_6_, 99.7%), urea (NH_2_CONH_2_, 99.5%), and uric acid (C_5_H_4_N_4_O_3_, 99%) were procured from Sigma-Aldrich (USA) unless otherwise stated. The thiol-modified IL-6 aptamer was purchased from Sigma-Aldrich (USA). Human IL-6 recombinant protein was obtained using the EZIL6 Human IL-6 ELISA Kit (Sigma-Aldrich, USA). 6-Mercapto-1-hexanol (MCH), used for electrode passivation, was also sourced from Sigma-Aldrich (USA). Artificial human serum (H4522, from human male AB plasma, USA origin, sterile-filtered) was obtained from Sigma-Aldrich (USA) and used as the real sample matrix for aptasensor validation. Additional interfering proteins, including tumor necrosis factor-alpha (TNF-α), ascorbic acid, urea, and uric acid, were used for selectivity studies. Gold screen-printed electrodes (Au-SPEs) were procured from a reputable supplier and used as received. Fresh phosphate-buffered saline (PBS, pH 7.5) was prepared by dissolving NaH_2_PO_4_ and NaH_2_PO_4_·2H_2_O in Milli-Q water and stored at 4 °C throughout the experiment. Milli-Q water was used throughout all experimental procedures. All reagents were of analytical grade and used without further purification.

### 2.2 Instrumentation

#### 2.2.1 Materials characterization

Atomic force microscopy (AFM) was performed using a Bio-AFM system (Bruker NanoWizard Sense AFM) to image surface topography and nanoscale features. For this, the samples were drop-casted onto the mica sheet and vacuum-dried. Dynamic light scattering (DLS) measurements were performed using a Nano ZS instrument (Malvern Instruments) to determine the hydrodynamic diameter and polydispersity index of the samples. For this, the samples were dispersed in deionized water (DI), and particle size was calculated using the Stokes-Einstein equation. The Zeta potential was also measured using the same instrument. Field-emission scanning electron microscopy (FESEM) was performed using a JSM-7600F (JEOL) to examine surface morphology and microstructure. Elemental analysis was performed using the attached energy-dispersive X-ray spectroscopy (EDX) system. Fourier transform infrared (FTIR) spectroscopy was performed using a PerkinElmer FTIR spectrometer to identify the functional groups. Spectra were recorded over the range of 4000-450 cm^-1^. The photoluminescence (PL) spectra were recorded using a Horiba Jobin Yvon Fluorolog-3 spectrophotometer. Raman spectroscopy was performed using a confocal Raman spectrometer (Renishaw, UK) to investigate vibrational properties. Spectra were recorded under ambient conditions using a laser excitation wavelength of 830 nm. For this, the samples were drop-cast onto the coverslips and vacuum-dried. Transmission electron microscopy (TEM) was conducted using an FEI Titan Themis 60-300 TEM equipped with an EDS detector and FEI Ceta 4k × 4k camera. TEM, high-resolution TEM (HRTEM), selected-area electron diffraction (SAED), and EDX were used to examine the morphology, crystallinity, and elemental composition of the samples. For this, the samples were dispersed in ethanol by ultrasonication. The dispersion was drop-cast onto a carbon-coated copper grid and dried overnight at RT. The thermogravimetric analysis (TGA) of the lyophilized samples was conducted with a TGA 4000 (PerkinElmer) under N2 flow from 50 to 780 °C at a heating rate of 10 °C min^-1^. UV-visible absorption spectra were recorded using a Thermo Scientific Orion AquaMate Vis and UV-Vis Spectrophotometer to analyze the optical properties of samples diluted in deionized water. Measurements were carried out over the 200-800 nm wavelength range for absorbance data collection. X-ray photoelectron spectroscopy (XPS) measurements were performed using a K-Alpha XPS system, equipped with a micro-focused Al Kα monochromatic X-ray source and a hemispherical analyzer, to investigate the surface elemental composition and chemical states. For this, the samples were drop-cast onto the coverslips and vacuum-dried. The phase structure and crystalline nature of the samples were investigated using grazing incidence X-ray diffraction (GIXRD) on a Bruker D8 DISCOVER powder X-ray diffractometer equipped with a Cu Kα radiation source. Data were collected in a 2θ scan mode over the angular range of 5°-90° (2θ). Cell viability was assessed by the MTT assay using a Bio-Rad 96-well plate reader at 562nm to quantify formazan formation spectrophotometrically. Confocal imaging was performed using a Leica TCS SP8 confocal laser scanning microscope to assess cell viability, membrane integrity, and oxidative stress at high spatial resolution. Wound-healing assays were monitored and imaged using a Nikon ECLIPSE Ts2 inverted microscope equipped with high-quality CFI60 optics and LED illumination for bright-field observation of cell migration.

#### 2.2.2 Electrochemical characterization

The fabricated aptasensor electrodes were electrochemically characterized using a PalmSens EmStat4S portable potentiostat or galvanostat connected to a standard three-electrode system. In this setup, the gold screen-printed electrode (Au-SPE) acted as the working electrode. The counter and reference electrodes were incorporated into the same SPE chip, eliminating the need for external electrode connections and creating a compact, reproducible measurement setup. All electrochemical data were collected and processed with PSTrace software. The measurements took place at room temperature under normal conditions. Cyclic voltammetry (CV) and differential pulse voltammetry (DPV) measurements were performed in PBS (pH 7.5) containing 5 mM [Fe(CN)_6_]^3-/4-^ as the redox probe. For CV measurements, the potential was scanned from -0.8 to +0.8 V at 50 mV s^-1^.

### 2.3 Synthesis of graphitic carbon nitride incorporated chitosan hydrogel

#### 2.3.1 Synthesis of g-C_3_N_4_

g-C_3_N_4_ was synthesized via thermal polymerization of urea. Briefly, 10g of urea was placed in a covered alumina crucible sealed with aluminum foil to create a semi-closed environment that facilitates condensation while minimizing material loss. The crucible was then transferred to a muffle furnace and heated to 550 °C at a controlled ramp rate of 3 °C/min, followed by calcination at this temperature for 4 hours and then cooled to room temperature. During this process, urea undergoes stepwise thermal decomposition and polycondensation, forming a layered graphitic carbon nitride framework that yields a pale-yellow solid. The obtained product was collected, thoroughly washed with ethanol and deionized water, centrifuged at 8000 rpm to remove residual impurities, and dried overnight at 80 °C. Finally, the dried material was ground into a fine powder using a mortar and pestle and stored in a sealed vial for further use^57,58^. **Scheme 1(a)** provides an illustrative depiction of the comprehensive synthesis process of g-C_3_N_4_.

#### 2.3.2 Synthesis of chitosan hydrogel

The chitosan hydrogel was prepared via chemical crosslinking with glutaraldehyde. Briefly, a 1% (w/v) chitosan solution was prepared by dissolving chitosan powder in 1% (v/v) acetic acid under continuous stirring, adding the powder gradually to ensure complete dissolution and prevent aggregation. The resulting viscous solution was then crosslinked by dropwise addition of glutaraldehyde at regular intervals (one drop every 2-3 min) under constant stirring as demonstrated in **Scheme 1(b)**. The crosslinking reaction occurs through the interaction of chitosan amine groups with glutaraldehyde aldehyde groups, forming covalent imine (Schiff base) linkages that stabilize the hydrogel network. The mixture progressively transformed into a gel, indicating the formation of a three-dimensional crosslinked chitosan hydrogel network^59^.

#### 2.3.3 Synthesis of g-C_3_N_4_@CS hydrogel

The g-C_3_N_4_@CS hydrogel was prepared by incorporating pre-synthesized g-C_3_N_4_ into the chitosan hydrogel matrix. Briefly, g-C_3_N_4_ powder was dispersed in deionized water and ultrasonicated for 2-3 h to exfoliate the layered structure into g-C_3_N_4_ nanosheets. Subsequently, an appropriate amount of the exfoliated g-C_3_N_4_ nanosheet suspension was added to the prepared chitosan solution, and the mixture was ultrasonicated for 1-2 h to ensure uniform dispersion of g-C_3_N_4_ nanosheets within the polymer matrix, as represented in **Scheme 1(b)**. Ultrasonication facilitates the exfoliation and homogeneous distribution of g-C_3_N_4_, preventing agglomeration and enhancing interfacial interaction with chitosan.

**Scheme 1:**
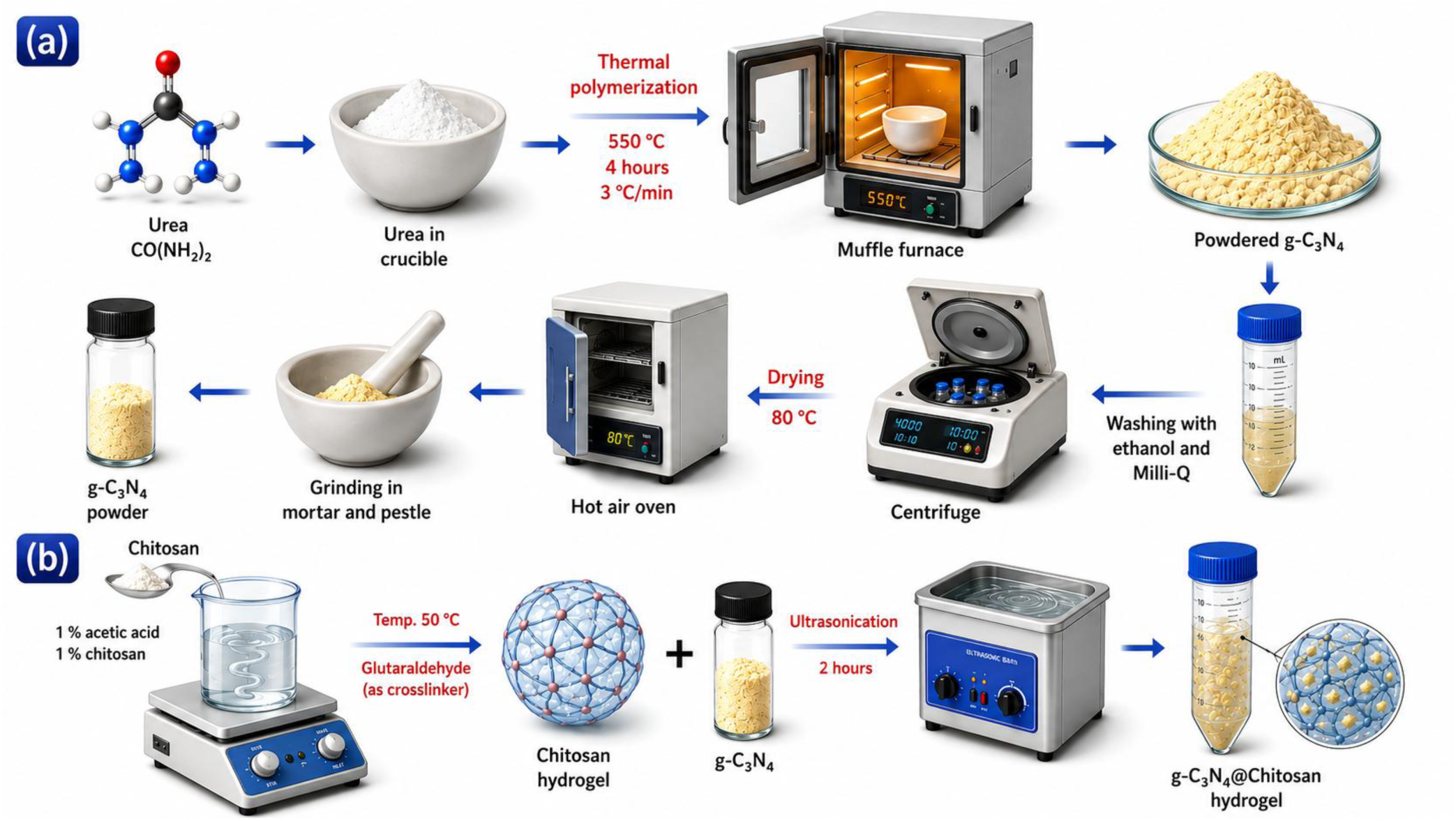
(a) Schematic illustration of g-C_3_N_4_ synthesis from urea via thermal polymerization (550 °C, 4 h), followed by washing, centrifugation, drying, and grinding to obtain powdered g-C_3_N_4_, and (b) Schematic representation of g-C_3_N_4_@CS hydrogel preparation involves forming a glutaraldehyde-crosslinked chitosan hydrogel followed by incorporation of g-C_3_N_4_ with ultrasonication.

### 2.4 Fabrication of the electrochemical aptasensor platform

#### 2.4.1 Aptamer preparation

The IL-6 aptamer sequence used in this study is shown in **Table S1**. The aptamer was modified with a thiol at the 5′ end to facilitate direct self-assembly onto the gold SPE surface. Before functionalizing the electrode, we reduced the thiolated aptamer to break any disulfide bonds and make sure free thiol groups were available. We mixed 1 μL of a 100 μM thiolated aptamer solution with 2 μL of a 25 mM tris(2-carboxyethyl)phosphine (TCEP) solution and incubated for 1 hour at room temperature. TCEP acts as a mild reducing agent that selectively cleaves disulfide linkages without altering the aptamer’s nucleotide structure. After the reduction, we diluted the aptamer to 47 μL in 1X PBS (pH 7.5) to achieve the desired working concentration for electrode functionalization.

#### 2.4.2 Electrode preparation

We used gold screen-printed electrodes (Au-SPEs) as the platform for our aptasensor. Before surface modification, we cleaned the Au-SPEs with isopropyl alcohol (IPA) for 2 minutes and rinsed them well with Milli-Q water. Next, we performed electrochemical cleaning by running 10 consecutive cyclic voltammetry scans in 0.1 M H_2_SO_4_ over a potential window of -0.8 to +0.8 V at a scan rate of 100 mV/s. This step removed any leftover surface contaminants and activated the gold surface. Then, we rinsed the electrodes again with Milli-Q water and dried them. Then, we drop-cast an optimized volume of the g-C_3_N_4_@CS hydrogel onto the cleaned Au-SPE surface and let it dry overnight at 25 °C, which produced the g-C_3_N_4_@CS/Au-SPE electrode.

#### 2.4.3 Development of the aptasensor platform

The g-C_3_N_4_@CS/Au-SPE electrode acted as the platform for building the label-free IL-6 aptasensor. We deposited a 5 μL aliquot of the reduced IL-6 aptamer solution (2 μM in 1X PBS, pH 7.5) onto the working electrode of the g-C_3_N_4_@CS/Au-SPE. We let it incubate overnight (about 18 hours) at room temperature. During this time, the free thiol groups at the 5′ end of the aptamer formed stable Au-S covalent bonds with the gold surface. The positively charged chitosan matrix in the g-C_3_N_4_@CS hydrogel provided additional electrostatic support for the negatively charged aptamer backbone, thereby ensuring stable, oriented aptamer immobilization. After incubation, we rinsed the electrode gently with Milli-Q water to remove any unbound aptamer strands, yielding the Apt-IL-6/g-C_3_N_4_@CS/Au-SPE electrode.

To cover the remaining bare gold spots not occupied by the aptamer and to reduce non-specific adsorption, we placed 5 μL of a 5 mM solution of 6-mercapto-1-hexanol (MCH) onto the Apt-IL-6/g-C3N4@CS/Au-SPE surface. We incubated it for 1 hour at room temperature. MCH serves as a short-chain alkanethiol blocking agent with dual purposes: it passivates the exposed gold surface by forming Au-S bonds. It helps orient the immobilized aptamer upright, thereby improving its accessibility to the IL-6 target. After passivation, we rinsed the electrode thoroughly with Milli-Q water to remove any loosely bound MCH molecules. We stored the resulting MCH/Apt-IL-6/g-C_3_N_4_@CS/Au-SPE aptasensor at 4 °C when not in use. This aptasensor was then used for all electrochemical characterization and IL-6 detection studies. A schematic of the entire fabrication process for the MCH/Apt-IL-6/g-C_3_N_4_@CS/Au-SPE aptasensor platform is shown in **Scheme 2**.

**Scheme 2:**
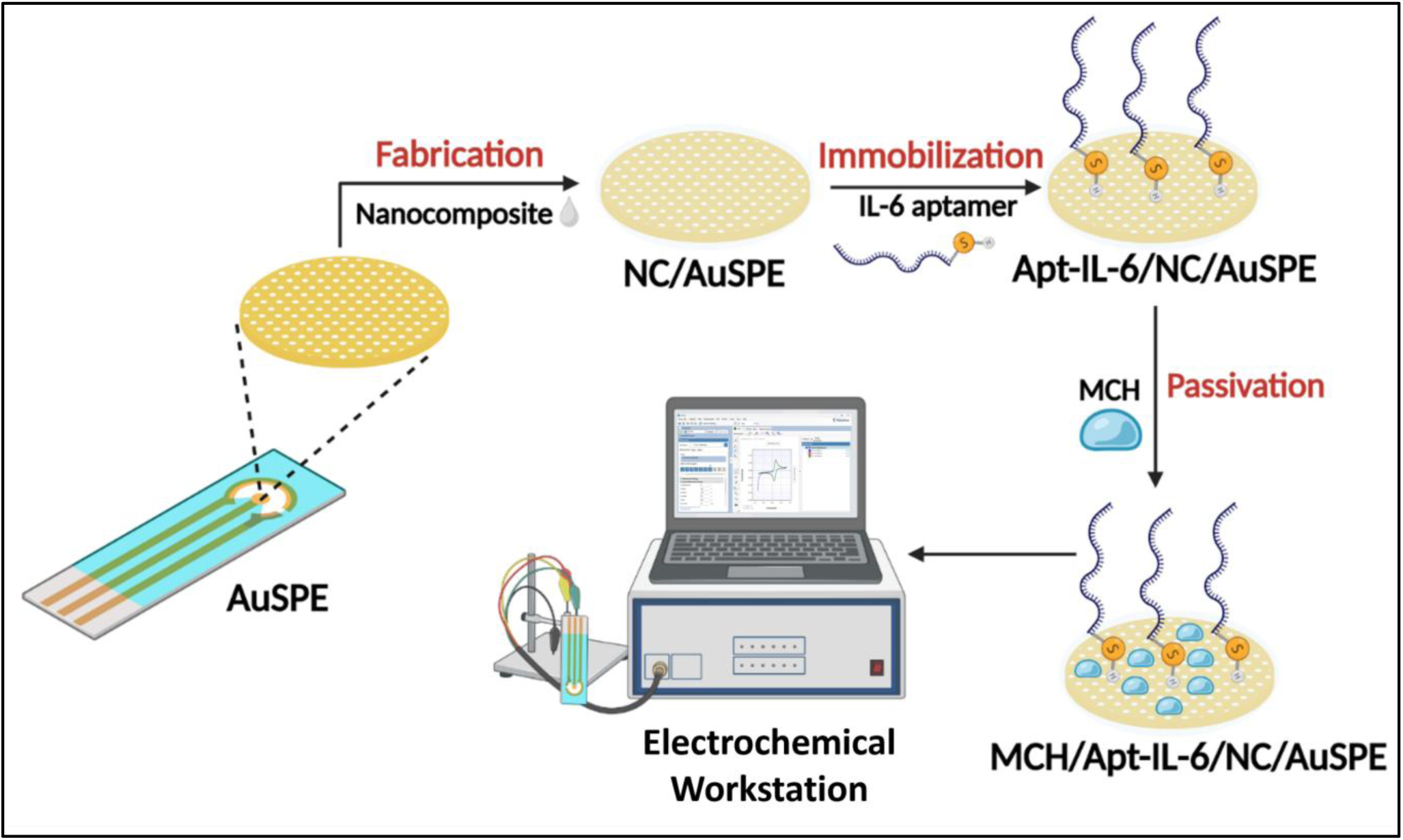
Schematic illustration of aptasensor fabrication on Au-SPE via g-C_3_N_4_@CS hydrogel deposition, IL-6 aptamer immobilization, MCH passivation, and electrochemical detection.

### 2.5 Analyte (IL-6) Sample Preparation

We prepared various concentrations of IL-6 protein by serial dilution in PBS (pH 7.5). Then, we applied these samples to the surface of the MCH/Apt-IL-6/g-C_3_N_4_@CS/Au-SPE aptaelectrode to achieve optimal response time. The IL-6 concentrations we examined covered a broad detection range to determine the linear dynamic range of the aptasensor. After the aptamer-protein binding interaction, we evaluated the resulting electrochemical responses using CV and DPV techniques in PBS (pH 7.5) containing 5 mM [Fe(CN)_6_]^3-/4-^ as the redox probe. For the DPV analysis, we set the pulse time and pulse potential at 0.02 s and 0.02 V, respectively. We conducted measurements at a scan rate of 0.05 V/s. The CV measurements covered a potential range of -0.8 to +0.8 V at a scan rate of 50 mV/s.

### 2.6 Preparation of Real Samples

To test the practical use of the MCH/Apt-IL-6/g-C_3_N_4_@CS/Au-SPE aptasensor in real clinical settings, we used artificial human serum as the sample matrix, as it mimics the composition of real human serum while ensuring reproducibility and safety in our experiments. We then spiked the supernatant with known concentrations of IL-6 protein across the detection range of the aptasensor, without further dilution. This step allowed us to evaluate the sensor’s recovery performance in a complex matrix. We used DPV to assess the extent of IL-6 binding to the MCH/Apt-IL-6/g-C_3_N_4_@CS/Au-SPE electrode. We calculated the percentage recovery to confirm the sensor’s accuracy and precision in analyzing real samples.

### 2.7 Biocompatibility Evaluation

The biocompatibility of the g-C_3_N_4_@CS hydrogel was assessed using an in vitro cell viability assay with RPE-1 cells cultured under standard conditions. Cells were exposed to sterilized samples for varying time durations, depending on the assay. We determined viability using MTT and confocal analysis, with untreated cells serving as the control. We performed both quantitative and qualitative analyses using absorbance measurements and fluorescence imaging. All experiments were done in triplicate.

## 3. RESULTS AND DISCUSSION

### 3.1 Structural Studies

The XRD patterns of g-C_3_N_4_, CS, and g-C_3_N_4_@CS hydrogel are shown in **Fig. 1(a)**. The pure chitosan showed a broad diffraction peak centered at 2θ = 10°. This indicates its semi-crystalline nature, arising from hydrogen bonding within the polymer chains. In contrast, the g-C_3_N_4_@CS hydrogel displayed two distinct diffraction peaks at about 13° and 27.4°. These correspond to the (100) in-plane structure of tri-s-triazine units and the (002) interlayer stacking of the conjugated aromatic layers of g-C_3_N_4_. The presence of the strong (002) peak confirms that g-C_3_N_4_ has been successfully added to the chitosan matrix. Additionally, the decrease in intensity and broadening of the chitosan peak in the hydrogel show a strong interaction between chitosan and g-C_3_N_4_. This results in partial loss of crystallinity and the formation of a uniform hydrogel.

**Figure 1:**
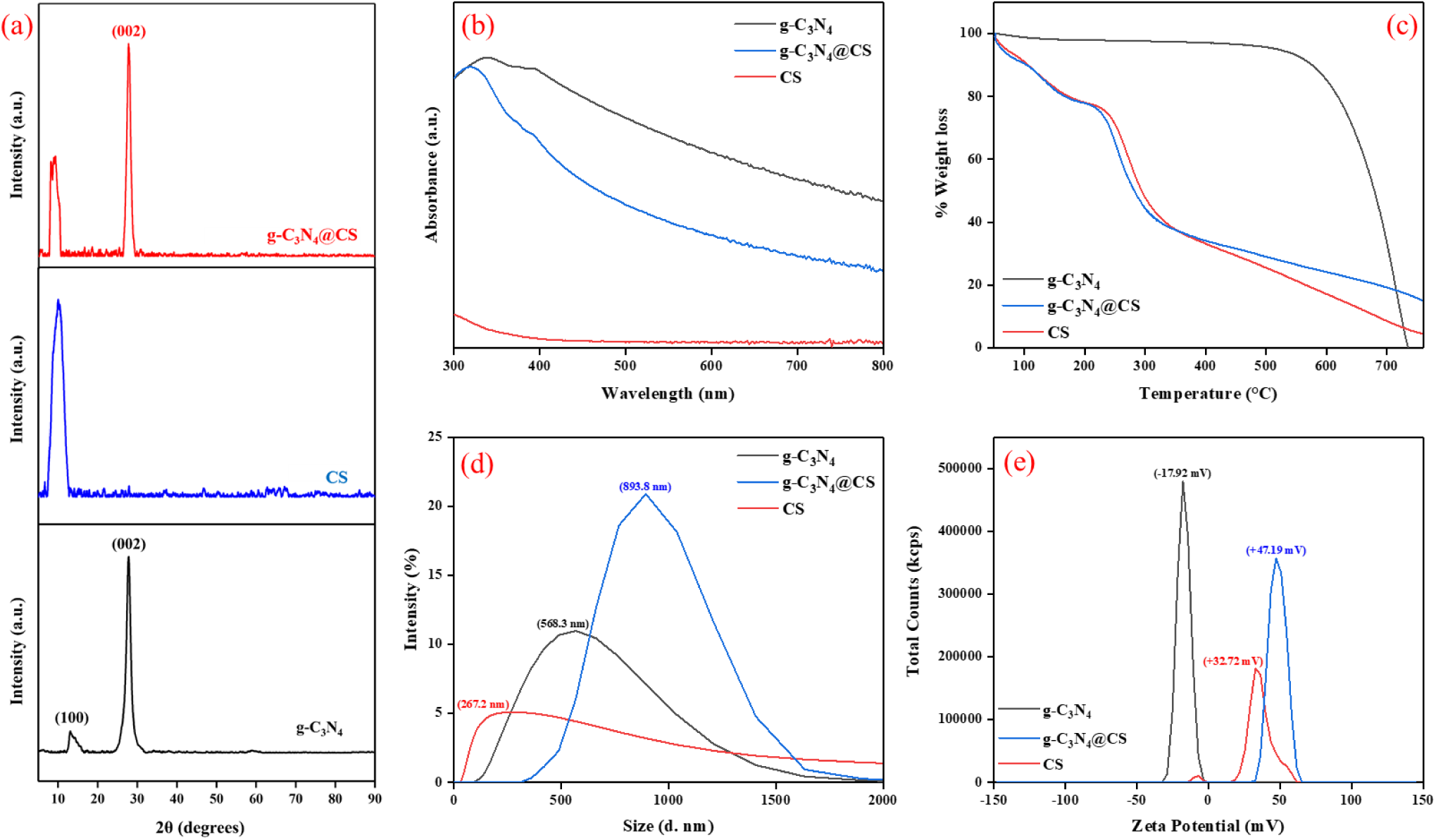
(a) XRD patterns, (b) UV-Vis spectra, (c) TGA curve, (d) DLS analysis, and (e) Zeta potential of g-C_3_N_4_, chitosan, and g-C_3_N_4_@CS hydrogel

The UV-visible absorption spectra of g-C_3_N_4_, chitosan, and the g-C_3_N_4_@CS hydrogel are shown in **Fig. 1(b)**. Pure g-C_3_N_4_ exhibits strong UV-visible absorption with an absorption edge around 400-450 nm. This is due to π-π* electronic transitions of the conjugated aromatic system. In comparison, chitosan has weak absorption only in the UV region because it lacks a conjugated system. The g-C_3_N_4_@CS hydrogel shows broadened absorption with a slightly shifted absorption edge compared to pure g-C_3_N_4_. This indicates electronic interaction between the g-C_3_N_4_ nanosheets and the chitosan matrix. The reduced intensity and wider absorption range suggest changes in the optical properties and a possible decrease in band gap energy.

Thermogravimetric analysis (TGA) was used to assess the thermal stability of g-C_3_N_4_, chitosan, and the g-C_3_N_4_@CS hydrogel **[Fig. 1(c)]**. Pure chitosan shows an initial weight loss below about 150 °C due to moisture removal. This is followed by significant degradation between 200 and 350 °C, which corresponds to the breakdown of the polymer backbone. The pristine g-C_3_N_4_ exhibits high thermal stability, with a sharp weight loss at 600-750 °C, attributed to the decomposition of its carbon-nitride structure. The g-C_3_N_4_@CS hydrogel exhibits a multi-step weight-loss pattern. The initial weight loss below around 150 °C is due to moisture removal. The gradual weight loss between 250 and 330 °C relates to the breakdown of chitosan. The significant weight loss at higher temperatures confirms the decomposition of g-C_3_N_4_.

The DLS analysis shows that chitosan has a broad size distribution centered at approximately 267 nm [**Fig. 1(d)]**. This suggests that polymeric aggregates are present in the solution. Pure g-C_3_N_4_ has a larger hydrodynamic diameter of about 568 nm. This size is due to the aggregation of layered nanosheets. The g-C_3_N_4_@CS hydrogel has a particle size of around 893 nm, which is even larger than that. This confirms that a hydrogel structure forms through the interaction between chitosan chains and g-C_3_N_4_ sheets. The increase in size results from polymer coating and aggregation effects, leading to a larger hydrodynamic diameter. The broad peaks indicate a polydisperse system, a common feature of polymer-based hydrogels. The zeta potential analysis shows that g-C_3_N_4_ has a negative surface charge of about -18 mV. In contrast, chitosan has a strongly positive charge of approximately +47 mV due to protonated amino groups. The g-C_3_N_4_@CS hydrogel has an intermediate zeta potential of approximately +30 to +36 mV **[Fig. 1(e)]**. This indicates an electrostatic interaction between the components with opposite charges. The movement toward lower positive values confirms that the hydrogel has formed successfully and that some charge neutralization has occurred. Additionally, the zeta potential’s magnitude suggests that the hydrogel system has moderate colloidal stability.

FT-IR spectra of g-C_3_N_4_, chitosan, and the g-C_3_N_4_@CS hydrogel are shown in **Fig. S1**. The FT-IR spectrum of g-C_3_N_4_ shows characteristic absorption peaks between 1200 and 1650 cm^-1^. These bands correspond to stretching vibrations of aromatic C-N and C=N heterocycles. There is also a distinct peak around 810 cm^-1^ that is linked to the breathing mode of the tri-s-triazine units. A broad band in the range of 3000 to 3400 cm^-1^ is due to N-H stretching vibrations. The spectrum of pure chitosan shows a broad peak from 3200 to 3400 cm^-1^, corresponding to overlapping O-H and N-H stretching vibrations. The peaks at 2922 cm^-1^ and 2871 cm^-1^ represent C-H stretching vibrations. The band at 1643 cm^-1^ was attributed to C=N stretching associated with Schiff base formation after glutaraldehyde crosslinking, while the band at 1549 cm^-1^ corresponded to N-H bending of residual amino groups. The peaks from 1400 to 1000 cm^-1^ were assigned to CH_2_ bending and C-O-C/C-O stretching vibrations of the chitosan backbone, confirming the presence of the polysaccharide network. In the g-C_3_N_4_@CS hydrogel, we observe characteristic peaks from both g-C_3_N_4_ and chitosan, with minor shifts and broadening, particularly in the 3000-3400 cm^-1^ and 1200-1650 cm^-1^ regions. These changes indicate strong interactions between the amino groups of chitosan and the nitrogen-rich sites of g-C_3_N_4_, likely mediated through hydrogen bonding.

The XPS survey spectrum confirms the elemental composition of the g-C_3_N_4_@CS hydrogel, as shown in **Fig. 2(a)**. It shows characteristic peaks for C 1s at 287 eV, N 1s at 400 eV, and O 1s at 533 eV. The carbon and nitrogen come from the graphitic carbon nitride framework. The oxygen peak is due to hydroxyl and ether groups in chitosan as well as adsorbed species (H_2_O and CO_2_). The relatively strong O 1s signal further confirms the successful addition of chitosan into the hydrogel structure. The C 1s XPS spectrum was deconvoluted into three main peaks. The peak at 284.6 eV corresponds to C-C/C=C bonds, often associated with adventitious carbon. The peak at 286.2 eV is associated with C-O/C-N bonds that come from the chitosan component. The largest peak at 288.2 eV is due to sp^2^-hybridized carbon in N-C=N groups, which is a key feature of the graphitic carbon nitride structure **[Fig. 2(b)]**. The N 1s spectra show four peaks at 398.4 eV, 399.1 eV, 399.6 eV, and 400.6 eV. These correspond to the sp^2^ C-N=C bonds, C-NH-C bonds, bridge N in the N-(C)_3_ groups, and the sp^2^-bonded N in the aromatic heterocyclic **[Fig. 2(c)]**. The O 1s spectrum shows two distinct peaks at 530.8 eV and 532.6 eV. The peak at 530.8 eV is linked to carbonyl oxygen (C=O) species. The higher binding-energy peak at 532.6 eV corresponds to C-O and O-H groups, including hydroxyl functionalities and adsorbed water molecules on the surface **[Fig. 2(d)]**.

**Figure 2:**
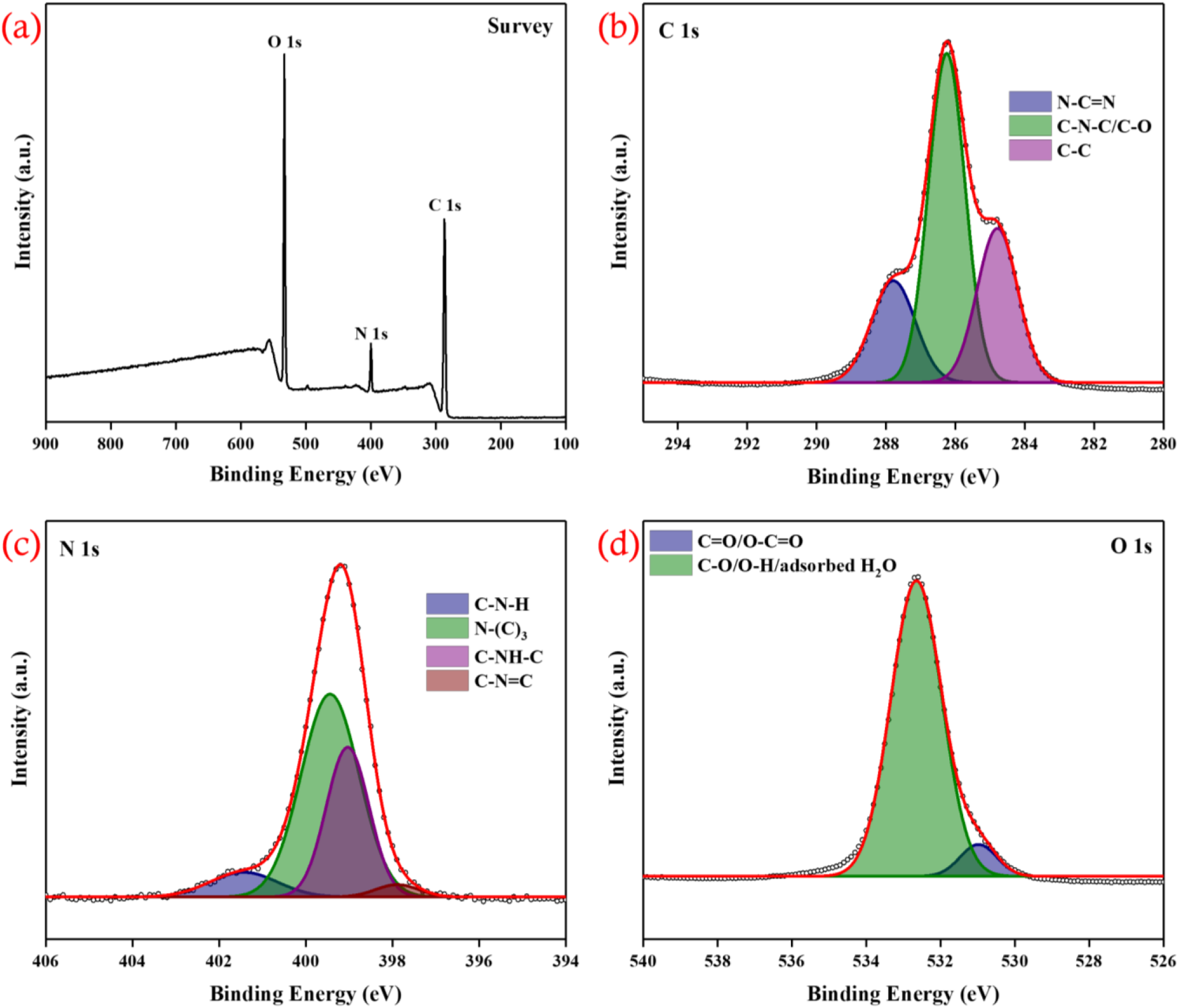
(a) XPS survey spectrum, (b) C 1s spectra, (c) N 1s spectra, and (d) O 1s spectra of g-C_3_N_4_@CS hydrogel

### 3.2 Morphological Studies

AFM characterization was performed to examine the surface structures of g-C_3_N_4_, chitosan hydrogel, and the g-C_3_N_4_@CS hydrogel films. As shown in **Fig. 3(a)**, pristine g-C_3_N_4_ had a rough, clustered flake-like appearance with a height variation of up to 916 nm, typical of its multilayered 2D sheet structure. The chitosan hydrogel **[Fig. 3(b)]** showed a very smooth and even surface with a height range of only 2.93 nm, confirming the creation of a continuous and uniform polymer film. In contrast, the g-C_3_N_4_@CS hydrogel **[Fig. 3(c)]** featured well-dispersed globular bumps spread across a continuous matrix with an intermediate height range of 366 nm. This indicates successful intercalation and stabilization of g-C_3_N_4_ nanosheets within the chitosan network. The reduced clustering of g-C_3_N_4_ in the hydrogel, along with the increased surface roughness compared to pristine chitosan, shows that the hydrogel offers a suitable electroactive interface with a larger surface area for IL-6 aptamer attachment on the Au-SPE surface. The surface structure of g-C_3_N_4_, CS hydrogel, and the g-C_3_N_4_@CS hydrogel was examined using FE-SEM. Pristine g-C_3_N_4_ **[Fig. 3(d)]** showed a heavily clustered, crumpled flake-like structure with a typical layered nanosheet stacking, which matches its 2D multilayered design. The chitosan hydrogel **[Fig. 3(e)]** had a highly porous, interconnected 3D network with smooth pore walls. This reflects the well-formed scaffolding resulting from chitosan chain crosslinking. The FESEM image of the g-C_3_N_4_@CS hydrogel **[Fig. 3(f)]** shows a porous network structure that maintains the general interconnected scaffold of the chitosan hydrogel but has thicker pore walls, smaller pore sizes, and a rougher surface texture compared to pure chitosan. This change in structure is due to the strong electrostatic attraction between the negatively charged g-C_3_N_4_ nanosheets. These nanosheets have surface nitrogen lone pairs and oxygen functional groups that give them a negative zeta potential. In contrast, the chitosan matrix carries a positive zeta potential under working conditions because its protonated amine (–NH_3_^+^) groups are positively charged.

**Figure 3:**
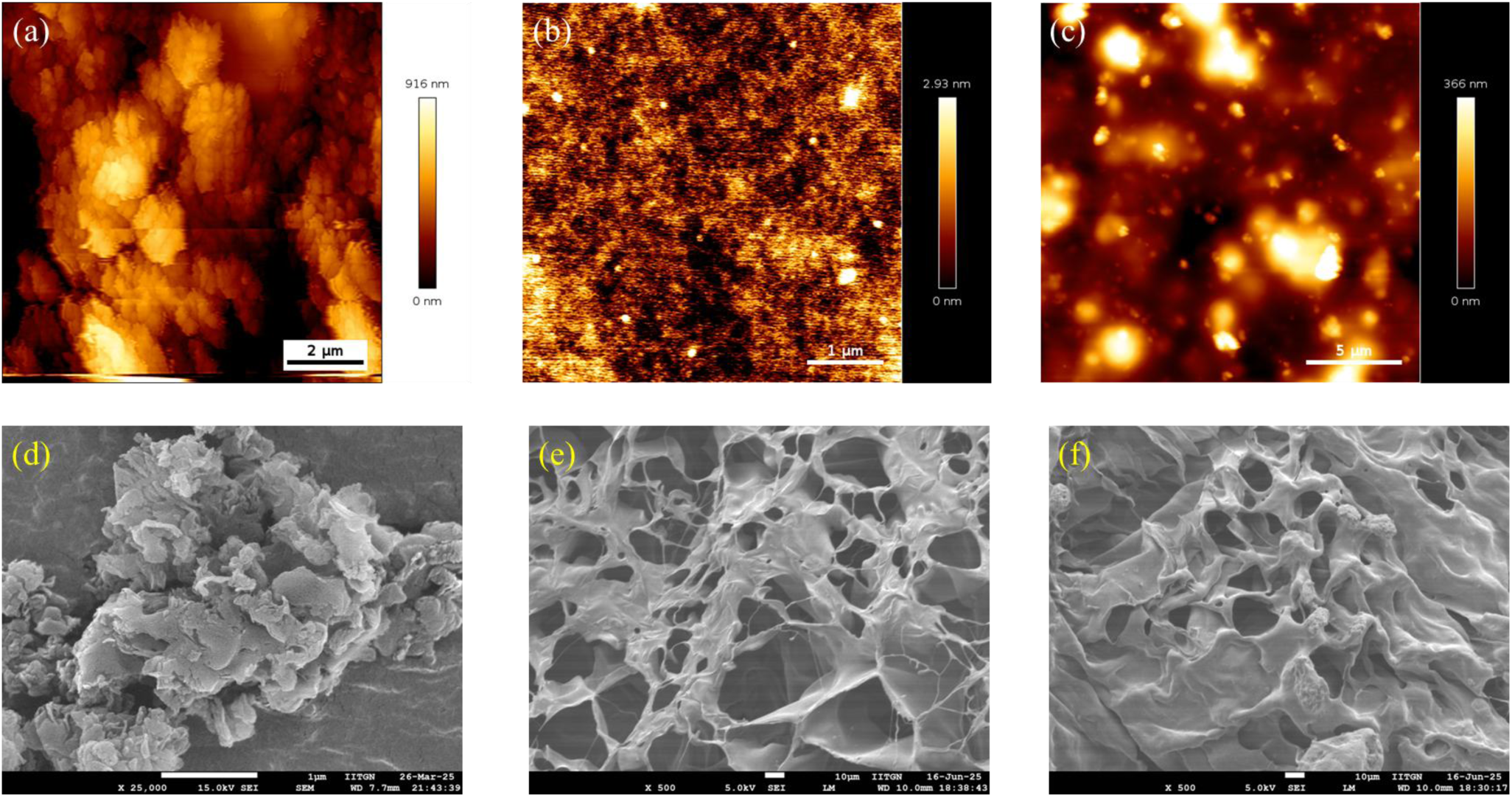
AFM image of (a) g-C_3_N_4_ (scale bar: 2 µm), (b) chitosan hydrogel (scale bar: 1 µm), and (c) g-C_3_N_4_@CS hydrogel (scale bar: 5 µm); and FESEM images of (d) g-C_3_N_4_ (scale bar: 1 µm), (e) chitosan hydrogel (scale bar: 10 µm), and (f) g-C_3_N_4_@CS hydrogel (scale bar: 5 µm).

Further, TEM, SAED, and HRTEM analyses were conducted to examine the nanostructural features of pristine g-C_3_N_4_ and the g-C_3_N_4_@CS hydrogel. The TEM image of pristine g-C_3_N_4_ **[Fig. 4(a)]** showed large, heavily aggregated nanosheet clusters with distinct folded and wrinkled sheet shapes, which matched its multilayered 2D structure. The corresponding SAED pattern **[Fig. 4(b)]** displayed a broad diffuse halo, confirming the semi-amorphous nature of the synthesized g-C_3_N_4_ with limited long-range crystalline ordering. HRTEM **[Fig. 4(c)]** revealed dense lattice fringes that matched the interlayer d-spacing of stacked g-C_3_N_4_ sheets. In contrast, the TEM image of the g-C_3_N_4_@CS hydrogel **[Fig. 4(d)]** showed a notably different morphology, with well-dispersed, significantly smaller g-C_3_N_4_ particles evenly distributed within the transparent chitosan matrix. This change confirmed effective disaggregation of g-C_3_N_4_, which resulted from the electrostatic attraction between the negatively charged g-C_3_N_4_ surface and the positively charged chitosan chains. The SAED pattern of the g-C_3_N_4_@CS hydrogel **[Fig. 4(e)]** exhibited a broader diffuse halo compared to pristine g-C_3_N_4_. This was due to the smaller g-C_3_N_4_ domain size and the additional amorphous contribution from the chitosan phase, with no new crystalline reflections observed. This finding confirmed that hydrogel formation occurs through physical electrostatic interaction rather than chemical bonding. The HRTEM image **[Fig. 4(f)]** showed significantly reduced lattice fringe contrast compared to pristine g-C_3_N_4_, consistent with chitosan encapsulating the g-C_3_N_4_ nanosheets and a decrease in effective layer stacking. Overall, these observations confirm the successful formation of a homogeneous g-C_3_N_4_@CS hydrogel, in which electrostatic complexation promotes uniform dispersion of g-C_3_N_4_ and prevents re-aggregation, creating a structurally favorable interface for IL-6 aptasensor fabrication.

**Figure 4:**
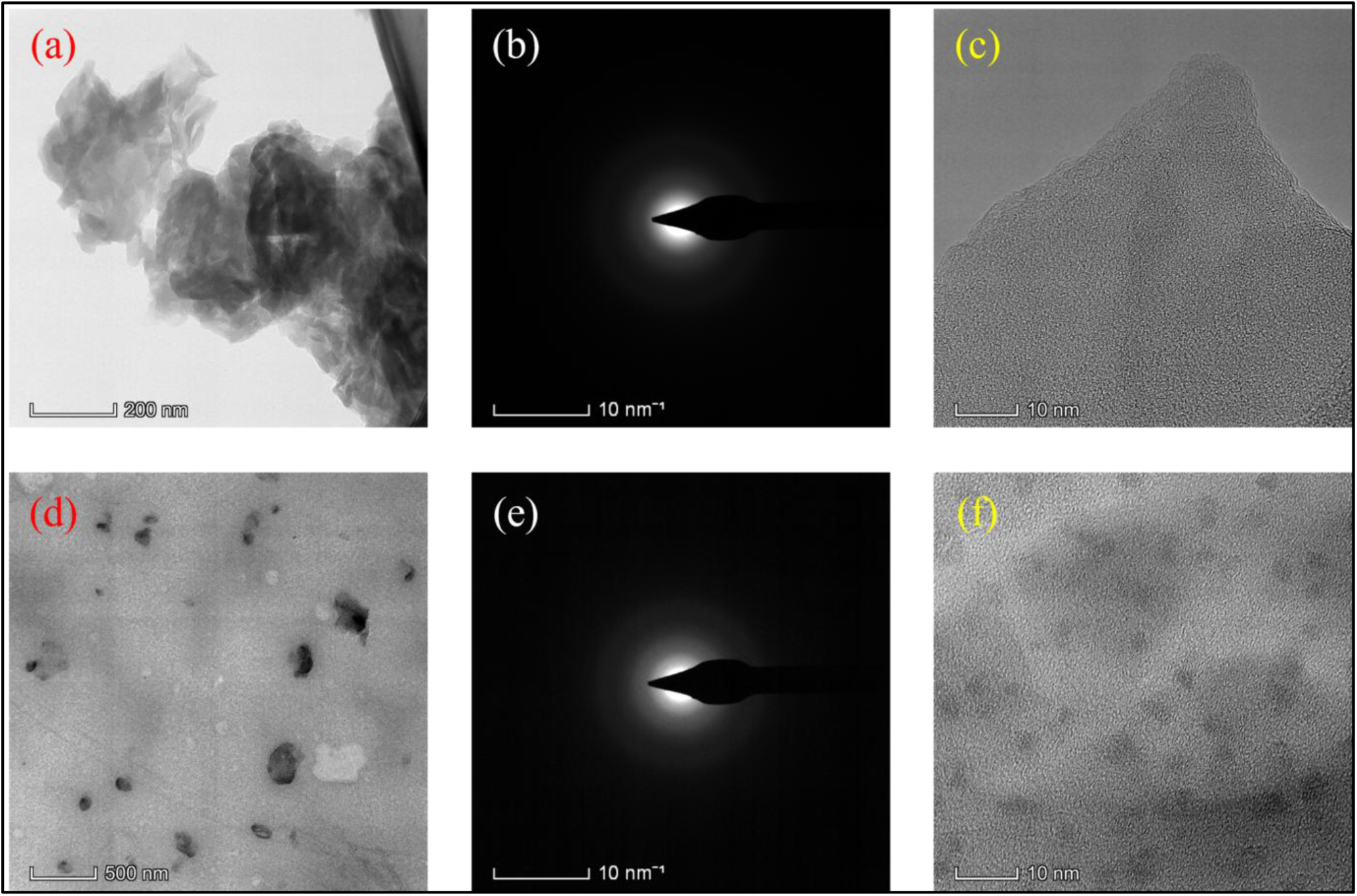
(a) TEM image, (b) SAED pattern, and (c) HR-TEM image of g-C_3_N_4_; (d) TEM image, (e) SAED pattern, and (f) HR-TEM image of g-C_3_N_4_@CS hydrogel.

### 3.3 Biocompatibility Studies

The biocompatibility of the g-C_3_N_4_@CS hydrogel was evaluated at 1, 5, and 10 mg/mL using the MTT cell viability assay **(Fig. 5(a))** and confocal fluorescence microscopy **(Fig. 5(b & c)**. The MTT results showed that higher concentrations were cytotoxic. At 1 and 5 mg/mL, the g-C_3_N_4_@CS hydrogel showed acceptable cell viability, similar to that of the chitosan hydrogel control. However, at 10 mg/mL, cell viability decreased significantly, indicating cytotoxicity at this higher concentration. Confocal microscopy, which used DAPI for nuclear staining, MitoTracker for mitochondrial integrity, and Phalloidin for F-actin cytoskeleton staining, supported these results. Cells treated with g-C_3_N_4_@CS hydrogel at 5 mg/mL had a healthy, spread-out shape, an intact actin structure, clear nuclear integrity, and an active mitochondrial distribution, similar to the untreated control. This confirmed good cytocompatibility at this concentration. In contrast, cells exposed to g-C_3_N_4_@CS hydrogel at 10 mg/mL showed signs of stress, including condensed cytoskeleton, decreased spreading, and altered mitochondrial distribution. Based on these biocompatibility tests, 5 mg/mL was selected as the optimal concentration for modifying the Au-SPE hydrogel. This ensures enough electrochemical activity and good cytocompatibility for possible biomedical use.

**Figure 5:**
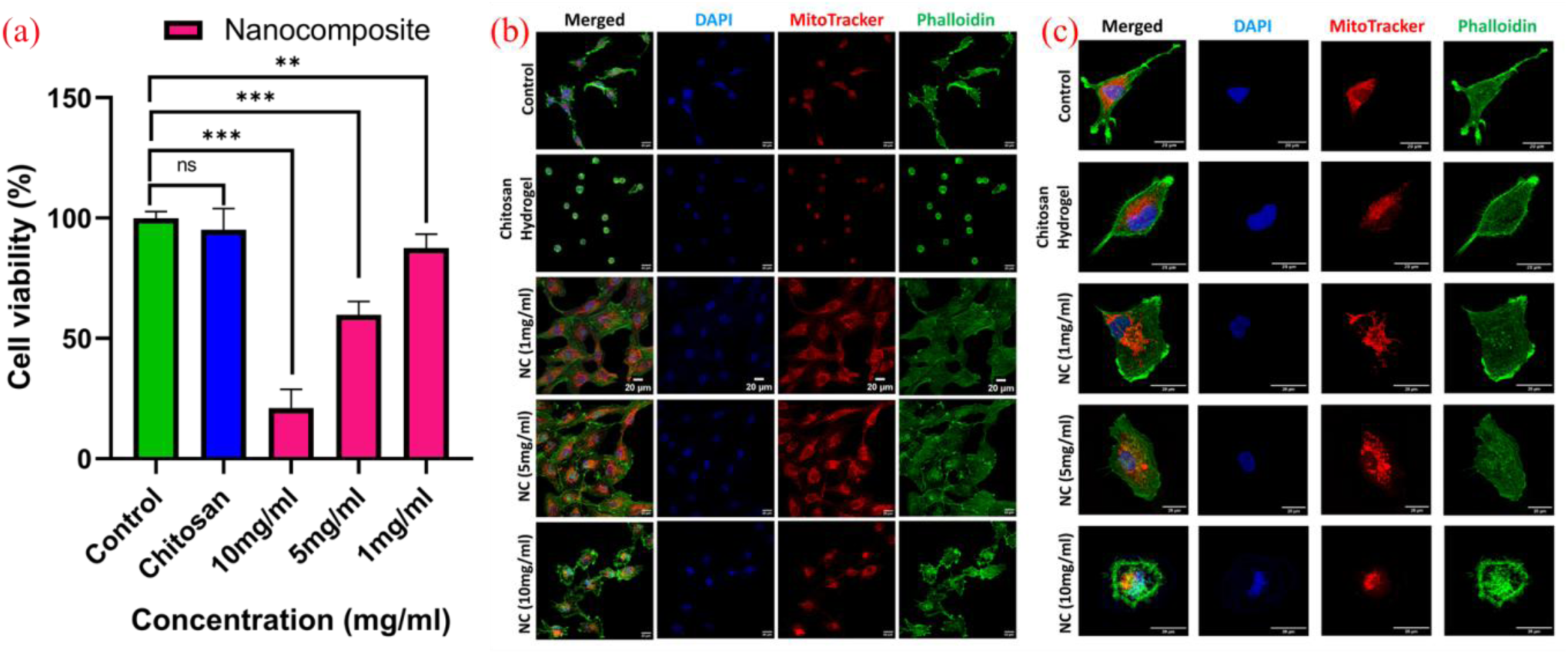
**(a)** MTT cell viability assessment of the g-C_3_N_4_@CS hydrogel at varying concentrations (1, 5, and 10 mg/mL) in comparison with untreated control and chitosan hydrogel groups. Cell viability is expressed as a percentage relative to the untreated control (n = 3, mean ± SD); **(b) & (c)** Confocal fluorescence microscopy images of cells treated with chitosan hydrogel and g-C_3_N_4_@CS hydrogel at concentrations of 1, 5, and 10 mg/mL in comparison with the untreated control group. Cells were co-stained with DAPI (blue, nuclear staining), MitoTracker (red, mitochondrial integrity), and Phalloidin (green, F-actin cytoskeletal morphology). Merged images represent the overlay of all three fluorescence channels. Scale bar = 20 μm

### 3.4 Electrochemical Studies

Electrochemical investigations were performed using a three-electrode setup that included an Ag/AgCl reference electrode (RE), a gold counter electrode (CE), and a gold screen-printed electrode (SPE) as the working electrode (WE). Using cyclic voltammetry (CV), we examined the electrochemical properties of the g-C_3_N_4_@CS hydrogel-modified AuSPEs with [Fe(CN)_6_]^3-/4-^ at each stage of surface modification. Although the kinetics and peak currents changed throughout the modification steps, the CVs of each electrode displayed a typical Fe^2+^/Fe^3+^ redox pair. The oxidation process (Fe^2+^ to Fe^3+^) showed a significantly lower peak current than the reduction process (Fe^2+^ to Fe^3+^). This suggests that the g-C_3_N_4_@CS hydrogel is leaning toward irreversibility and creating some electron-transfer resistance at the electrode surface. The reduction process is faster than the oxidation process due to the interfacial properties introduced by the g-C_3_N_4_@CS hydrogel. This behavior is likely due to the carbon nitride components acting as reducing agents. The nitrogen lone pairs and surface oxygen functional groups provide additional negative charge, which favors reduction over oxidation. Chitosan acts as a binder to stabilize g-C_3_N_4_ on the gold SPE surface and as an immobilization matrix. It further influences electron-transfer kinetics through its amine (-NH_2_) and hydroxyl (-OH) functional groups, thereby affecting the overall charge distribution at the interface.

#### 3.4.1 Optimization of experimental conditions

Optimizing the various experimental parameters is essential for developing a biosensor that is sensitive, repeatable, and reproducible. We adjusted various experimental conditions, starting with determining the optimal parameters for the g-C_3_N_4_@CS hydrogel modification. This included its concentration and the drop-casting volume applied to the Au-SPE surface. Other important factors we looked at were the volume of the IL-6 aptamer used for immobilization, the volume of MCH used to block nonspecific binding sites, and the aptamer-target interaction time, which corresponds to the response time. All of these factors directly affect the sensitivity and specificity of the aptasensor. We systematically optimized these parameters before evaluating the aptasensor’s performance to find the best experimental conditions. The optimized parameters are shown in **Fig. S2(A-C)**. The optimized values for each experimental parameter, along with the tested and selected values, are shown in **Table S2**.

#### 3.4.2 Effect of scan rate studies

The scan rate is crucial for the electrocatalytic behavior of any electrochemical biosensor. It directly affects charge transfer dynamics at the electrode interface. To better understand the electrochemical behavior of the aptasensor we created, we recorded CVs for the g-C_3_N_4_@CS/Au-SPE and MCH/Apt-IL-6/g-C_3_N_4_@CS/Au-SPE electrodes as the scan rate was gradually increased from 10 to 100 mV/s, as shown in **Fig. 6(A) and 6(B)**. In both cases, the anodic and cathodic peak currents increased with increasing scan rate, while the peak potentials remained nearly constant. This suggests that the electrode process is electrochemically reversible within the examined scan rate range. The lack of a significant peak potential shift at both the g-C_3_N_4_@CS/Au-SPE and MCH/Apt-IL-6/g-C_3_N_4_@CS/Au-SPE surfaces indicates that the electron-transfer kinetics at the modified electrode interface are fast enough to maintain reversibility as the scan rate increases. This behavior suggests that the g-C_3_N_4_@CS hydrogel, when attached to the gold SPE, enables efficient electron transfer without significant kinetic barriers. We confirmed that the charge-transfer mechanism is diffusion-controlled. This is shown by the linear relationship between the anodic peak current and the square root of the scan rate, a classic sign of diffusion-limited electrochemical processes. The preservation of peak reversibility across all scan rates underscores the favorable electron-transfer properties of the g-C_3_N_4_@CS hydrogel interface, which is especially beneficial for ensuring stable, reliable aptasensor performance under diverse measurement conditions.

**Figure 6:**
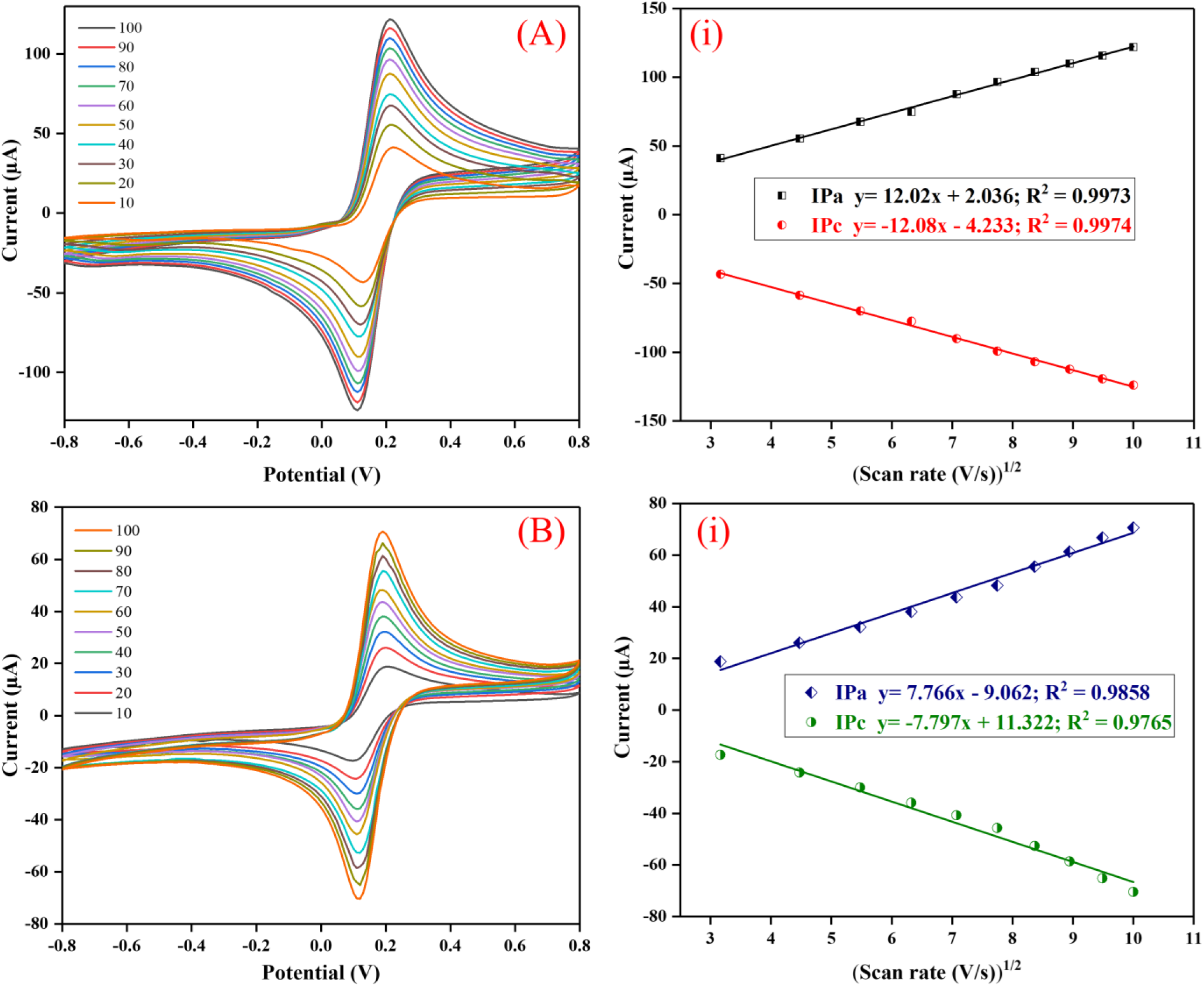
CV profiles of (A) the g-C_3_N_4_@CS/Au-SPE and (B) the MCH/Apt-IL-6/NC/Au-SPE aptaelectrode. A(i) and B(i) display the oxidation and reduction peak currents normalized to the square root of the scan rate.

**Fig. 6A(i) and 6B(i)** show the linear relationships between the oxidation peak current (Ipa) and the reduction peak current (Ipc) as a function of the square root of the scan rate (mV/s) and follow eq (1)-(4).

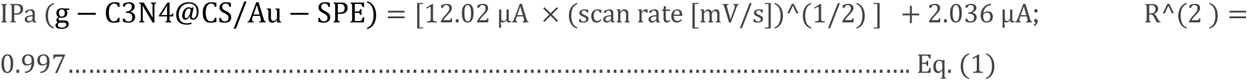

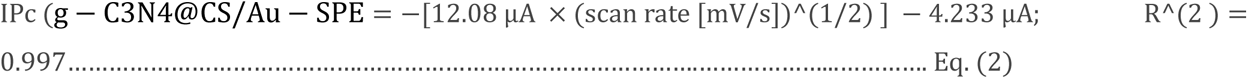

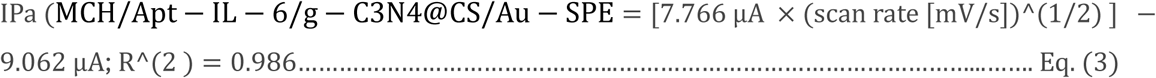

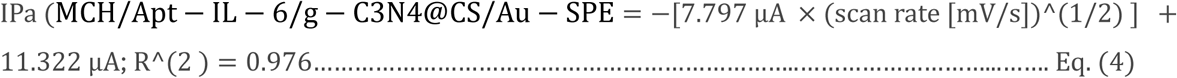

Additionally, there was no significant shift in the anodic or cathodic peak potentials at either electrode. The peak-to-peak separation (ΔE = Epa - Epc) remained largely consistent across all scan rates. This indicates that the electrochemical process at the modified electrode surface is fully reversible. It shows that electron-transfer kinetics at the g-C_3_N_4_@CS/Au-SPE interface are fast enough to maintain equilibrium even at higher scan rates. This reflects the effective charge transfer properties of the g-C_3_N_4_@CS hydrogel on the gold SPE surface.

The kinetic properties of the g-C_3_N_4_@CS/Au-SPE and the MCH/Apt-IL-6/g-C_3_N_4_@CS/Au-SPE electrode were thoroughly assessed to examine the electron transfer dynamics at each phase of aptasensor fabrication. **Table 1** presents the calculated electrochemical parameters for both electrodes, including cathodic peak current (Ipc), anodic peak current (Ipa), diffusion coefficient (D), average surface concentration (γ*), charge transfer rate constant (Ks), and electroactive surface area (Ae). A comparison of the diffusion coefficient (D) values between the g-C_3_N_4_@CS/Au-SPE and the MCH/Apt-IL-6/g-C_3_N_4_@CS/Au-SPE electrode shows that the latter had lower D values. This decrease in D is due to the sequential layering of biomolecules, specifically the IL-6 aptamer and the MCH blocking agent, which together act as physical and electrostatic barriers at the electrode surface.

**Table 1.**
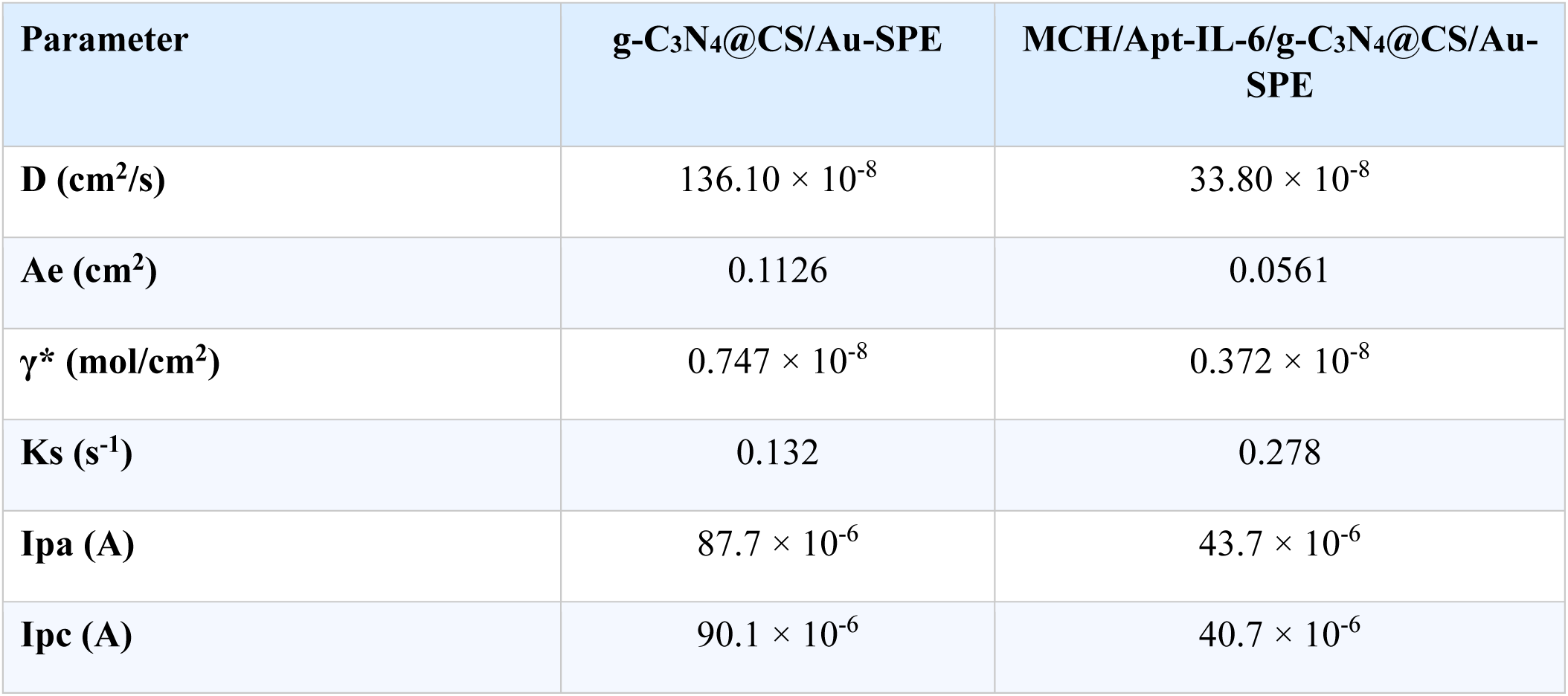
Predicted values of electrochemical parameters for electrodes.

| Parameter | g-C<sub>3</sub>N<sub>4</sub>@CS/Au-SPE | MCH/Apt-IL-6/g-C<sub>3</sub>N<sub>4</sub>@CS/Au-SPE |
| --- | --- | --- |
| **D (cm<sup>2</sup>/s)** | 136.10 × 10<sup>-8</sup> | 33.80 × 10<sup>-8</sup> |
| **Ae (cm<sup>2</sup>)** | 0.1126 | 0.0561 |
| **γ\* (mol/cm<sup>2</sup>)** | 0.747 × 10<sup>-8</sup> | 0.372 × 10<sup>-8</sup> |
| **Ks (s<sup>-1</sup>)** | 0.132 | 0.278 |
| **Ipa (A)** | 87.7 × 10<sup>-6</sup> | 43.7 × 10<sup>-6</sup> |
| **Ipc (A)** | 90.1 × 10<sup>-6</sup> | 40.7 × 10<sup>-6</sup> |

The aptamer, as a large nucleic acid strand, introduces steric hindrance, limiting access of the electroactive species ([Fe(CN)_6_]^3-/4-^) to the electrode surface. Moreover, the MCH monolayer helps orient the aptamer and blocks nonspecific adsorption sites, further limiting the diffusion of redox species to the gold SPE surface. These combined factors, steric hindrance, electrostatic repulsion, and surface passivation, reduce charge transfer efficiency and slow diffusion kinetics. This explains the lower D value for the MCH/aptamer/g-C_3_N_4_@CS/Au-SPE electrode compared to that of the bare g-C_3_N_4_@CS/Au-SPE electrode.

#### 3.4.3 pH studies

pH optimization studies aimed to find the best PBS buffer pH for the MCH/Apt-IL-6/g-C_3_N_4_@CS/Au-SPE aptaelectrode. The research used PBS with the [Fe(CN)_6_]^3-/4-^ redox couple at a constant scan rate of 50 mV s⁻¹. We evaluated the electrochemical response of the MCH/Apt-IL-6/g-C_3_N_4_@CS/Au-SPE aptaelectrode over a pH range of 6.0 to 8.0 using both CV and DPV. The results, shown in **Fig. 7(a)** and **(c)**, indicate a clear relationship between the electrochemical signal and buffer pH. This relationship arises from how pH affects the surface charge distribution of the IL-6 aptamer on the electrode, thereby altering the overall electrochemical response.

**Figure 7:**
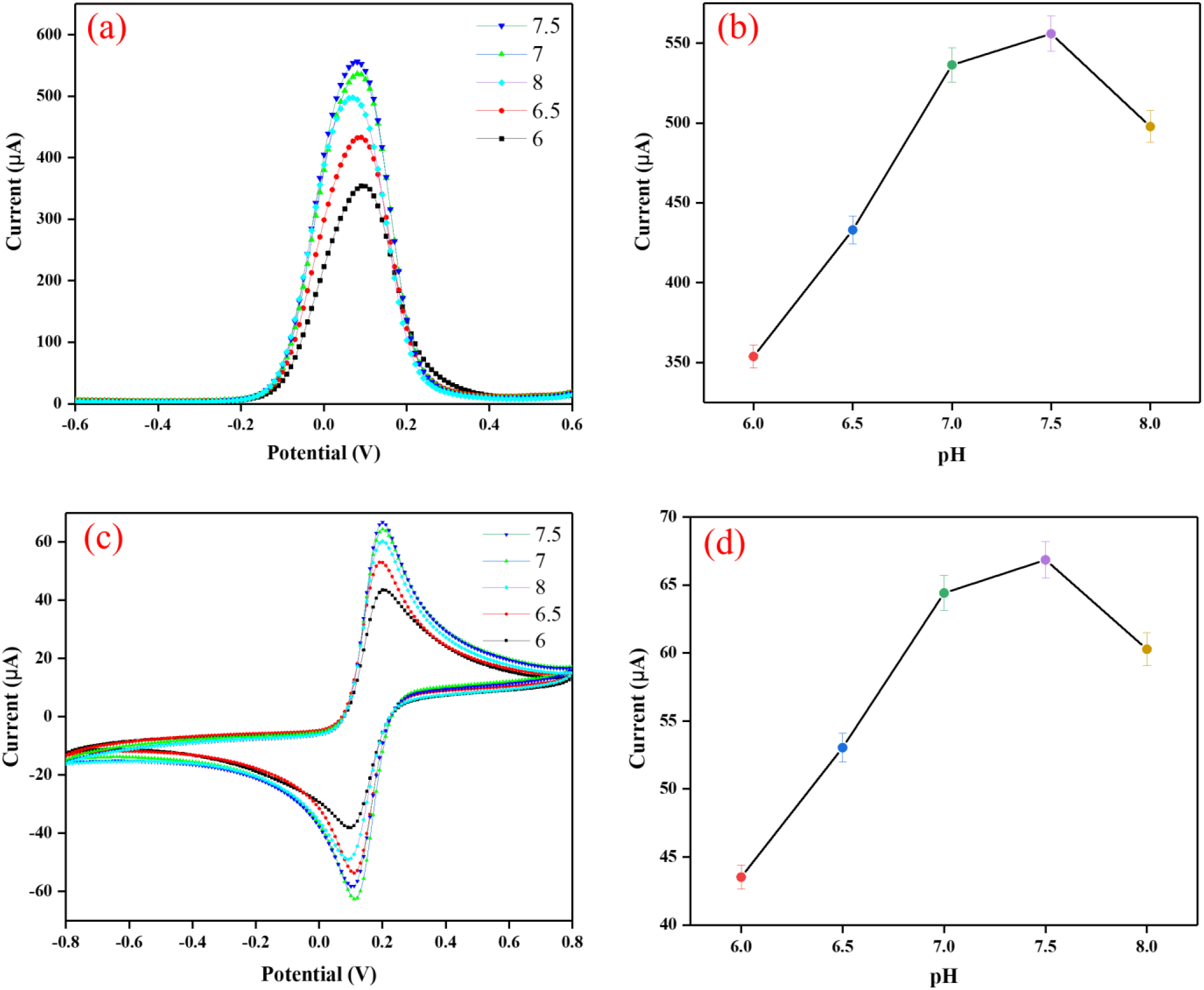
pH study using (a) CV and (c) DPV techniques with (b) and (d) the line plot demonstrating the correlation between the pH and the peak current (oxidation).

**Fig. 7(b)** and **(d)** show the oxidation peak current plotted against pH. The highest anodic current response occurred at pH 7.5. This slightly alkaline, nearly physiological pH provides the optimal environment for preserving the structure and activity of the immobilized IL-6 aptamer on the gold SPE surface. At pH below 7.5, partial protonation of the aptamer’s nucleotide residues can disrupt its optimal folding, leading to a weaker electrochemical signal. On the other hand, at pH values above 7.5, increased alkalinity leads to deprotonation and electrostatic repulsion in the aptamer backbone. This disrupts its three-dimensional shape and lowers the current response. The maximum response at pH 7.5 also fits well with the pH range typical in human serum and environments for detecting inflammatory biomarkers, making it especially appropriate for IL-6 sensing applications. Based on these findings, we selected pH 7.5 PBS as the optimal buffer for further electrochemical analysis and aptasensor performance assessment.

#### 3.4.4 Electrochemical studies of fabricated electrodes

The electron transport characteristics of the stepwise-fabricated aptasensor electrodes were investigated using both CV and DPV in PBS (pH 7.5) containing 5 mM [Fe(CN)_6_]^3-/4-^ as the redox probe. The DPV responses for the sequential electrode modifications are presented in **Fig. 8(c)**. The relative current comparison between the fabricated electrodes is shown in **Fig. 8(d)**. All CV measurements were conducted over a potential range of -0.8 V to +0.8 V at a scan rate of 50 mV/s [**Fig. 8(a) and (b)]**.

**Figure 8:**
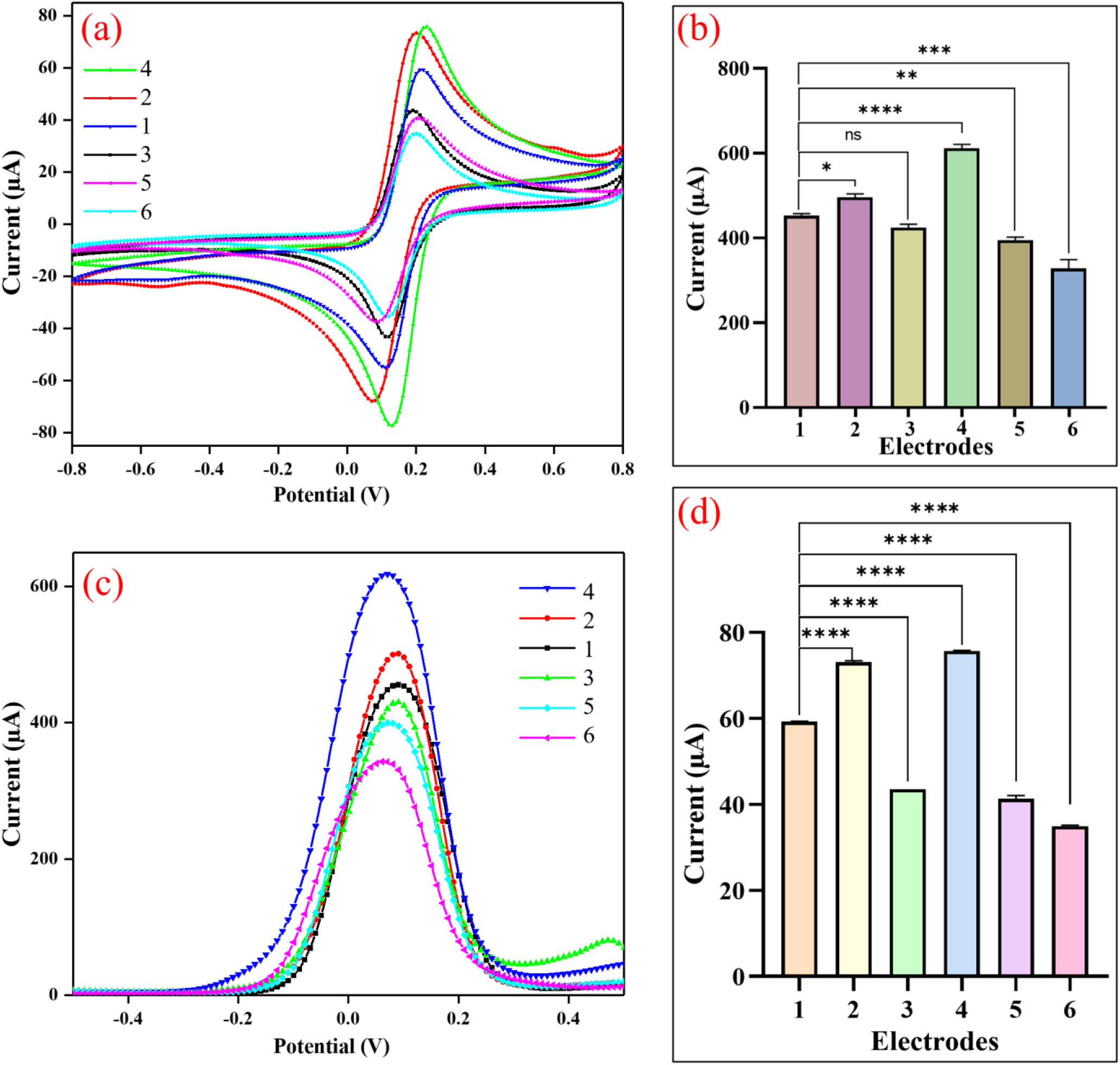
(a) CV and (b) DPV curves of various modified electrodes (1. Au-SPE, 2. g-C_3_N_4_/Au-SPE, 3. CS/Au-SPE, 4. g-C_3_N_4_@CS/Au-SPE, 5. Apt-IL-6/g-C_3_N_4_@CS/Au-SPE, and 6. MCH/Apt-IL-6/g-C_3_N_4_@CS/Au-SPE); (b) and (d) the bar plot between the peak current and various modified electrodes.

The bare Au-SPE exhibited a moderate peak current, reflecting the inherent electrochemical activity of the clean gold surface toward the [Fe(CN)_6_]^3-/4-^ redox couple. Upon drop-casting the g-C_3_N_4_@CS hydrogel onto the Au-SPE, the peak current increased appreciably, indicating enhanced electron transfer facilitated by the electrochemically active g-C_3_N_4_ component within the hydrogel. The improved electron transfer can be attributed to favorable electrostatic interactions between the hydrogel surface and [Fe(CN)_6_]^3-/4-^ ions, as well as to the increased electroactive surface area provided by the porous hydrogel architecture. Following the immobilization of the thiol-modified IL-6 aptamer onto the g-C_3_N_4_@CS/Au-SPE surface, a progressive decrease in peak current was observed alongside an increase in peak-to-peak separation (ΔEp), consistent with the introduction of a partially blocking biomolecular layer that sterically hinders the access of redox species to the electrode surface. The subsequent passivation with MCH further reduced the peak current, confirming the successful blocking of residual bare gold sites and non-specific adsorption regions. The insulating nature of the MCH monolayer, combined with its role in aptamer reorientation, collectively contributes to the additional attenuation of the electrochemical signal at the MCH/Apt-IL-6/g-C_3_N_4_@CS/Au-SPE aptaelectrode surface. These stepwise changes in current and corresponding ΔEp variations were similarly reflected in the DPV measurements, corroborating the CV findings and confirming the successful sequential fabrication of the aptasensor.

The electron transfer abilities of bare Au-SPE, g-C_3_N_4_@CS/Au-SPE, Apt-IL-6/g-C_3_N_4_@CS/Au-SPE, and MCH/Apt-IL-6/g-C_3_N_4_@CS/Au-SPE were also investigated in 0.2 M PBS containing 5 mM [Fe(CN)_6_]^3-/4-^ in the potential range between -0.8 and +0.8 V vs Ag/AgCl using CV at a scan rate of 50 mV/s. **Table 2** compares the measured oxidation and reduction peak current densities (Jpa and Jpc), anodic and cathodic peak potentials (Epa and Epc), and potential differences (ΔEp). Consequently, compared to other modified and bare Au-SPE electrodes, the MCH/Apt-IL-6/g-C_3_N_4_@CS/Au-SPE electrode showed higher ΔEp and more significant redox peak currents. This could be attributed to the favorable electrostatic interactions between the g-C_3_N_4_@CS surface and [Fe(CN)_6_]^3-/4-^ ions, along with the high electrical conductivity of g-C_3_N_4_@CS, which promotes rapid electron transfer. Moreover, the MCH blocking layer minimizes non-specific adsorption, while the successful immobilization of Apt-IL-6 further confirms the stepwise construction of the aptasensor platform for IL-6 detection.

**Table 2.**
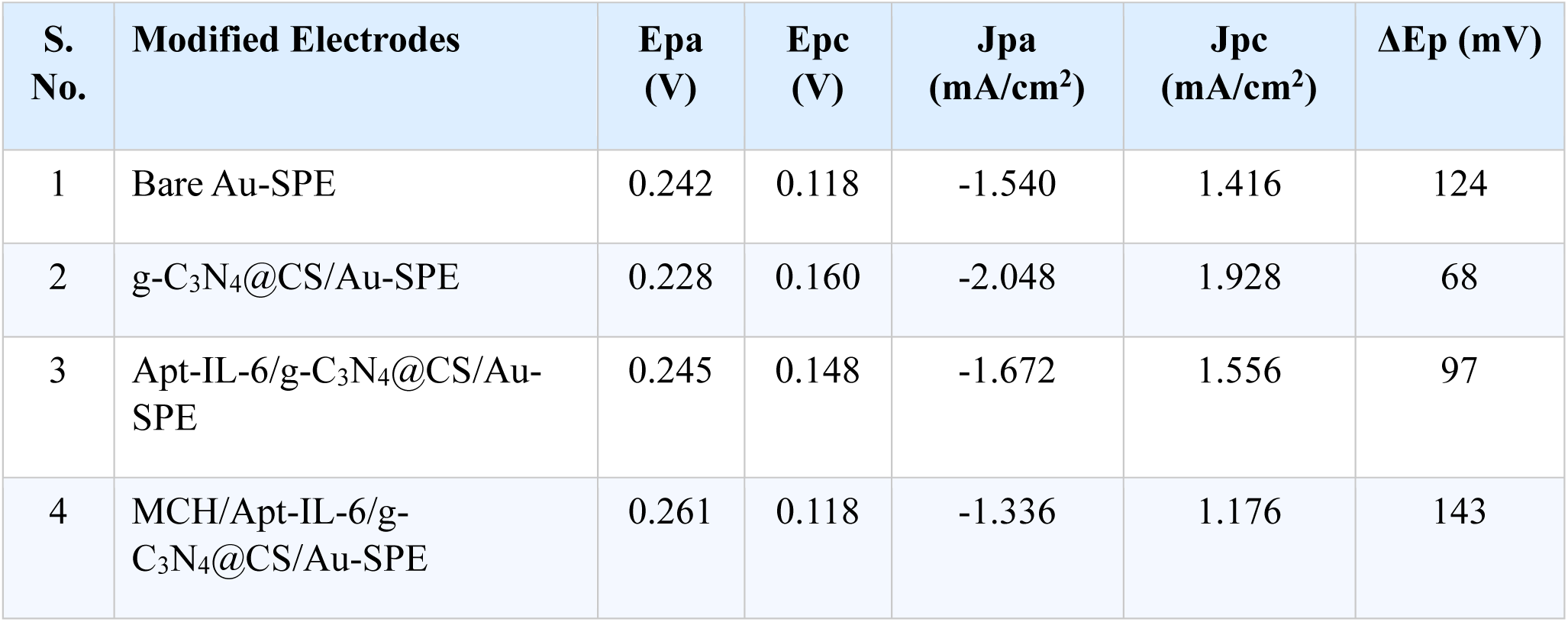
CV profiles of various modified electrodes and their potentials and current values in 0.2 M PBS containing redox species.

| S. No. | Modified Electrodes | E<sub>pa</sub> (V) | E<sub>pc</sub> (V) | J<sub>pa</sub> (mA/cm<sup>2</sup>) | J<sub>pc</sub> (mA/cm<sup>2</sup>) | ΔE<sub>p</sub> (mV) |
| --- | --- | --- | --- | --- | --- | --- |
| 1 | Bare Au-SPE | 0.242 | 0.118 | -1.540 | 1.416 | 124 |
| 2 | g-C<sub>3</sub>N<sub>4</sub>@CS/Au-SPE | 0.228 | 0.160 | -2.048 | 1.928 | 68 |
| 3 | Apt-IL-6/g-C<sub>3</sub>N<sub>4</sub>@CS/Au-SPE | 0.245 | 0.148 | -1.672 | 1.556 | 97 |
| 4 | MCH/Apt-IL-6/g-C<sub>3</sub>N<sub>4</sub>@CS/Au-SPE | 0.261 | 0.118 | -1.336 | 1.176 | 143 |

#### 3.4.5 Analytical Performance of the Aptasensor for IL-6 Detection

Before conducting electrochemical response investigations, a response-time optimization study was performed to determine the optimal incubation time required for IL-6 to interact with the MCH/Apt-IL-6/g-C_3_N_4_@CS/Au-SPE aptaelectrode surface. The current response was monitored at 1-minute intervals for up to 15 minutes, and an incubation time of 5 minutes was identified as optimal for achieving maximum, stable aptamer-protein binding. This optimized response time was subsequently applied to all electrochemical detection experiments.

DPV was employed as the primary analytical technique for IL-6 quantification owing to its superior sensitivity and effective suppression of non-faradaic background currents compared to CV. The electrochemical response of the MCH/Apt-IL-6/g-C_3_N_4_@CS/Au-SPE aptaelectrode was investigated across a wide range of IL-6 concentrations spanning from 1 fg/mL to 10000 pg/mL in PBS (pH 7.5) containing 5 mM [Fe(CN)_6_]^3-/4-^, over the potential range of -0.8 V to +0.8 V at a scan rate of 50 mV/s. As illustrated in **Fig. 9(a)**, the DPV peak current decreased progressively with increasing IL-6 concentration, following a signal-off response mechanism. This behavior arises from the specific binding of IL-6 to the immobilized aptamer, which forms a conformational aptamer-protein complex that acts as an electrically insulating barrier, progressively impeding the electron transfer between the aptaelectrode surface and the [Fe(CN)_6_]^3-/4-^ redox probe. The formation of this complex at higher IL-6 concentrations results in a thicker insulating layer on the MCH/Apt-IL-6/NC/Au-SPE surface, further restricting the accessibility of redox species and producing the observed concentration-dependent current attenuation. All measurements were performed in triplicate, and an enhanced view of the electrochemical response is provided in the inset of **Fig. 9(a)**.

**Figure 9:**
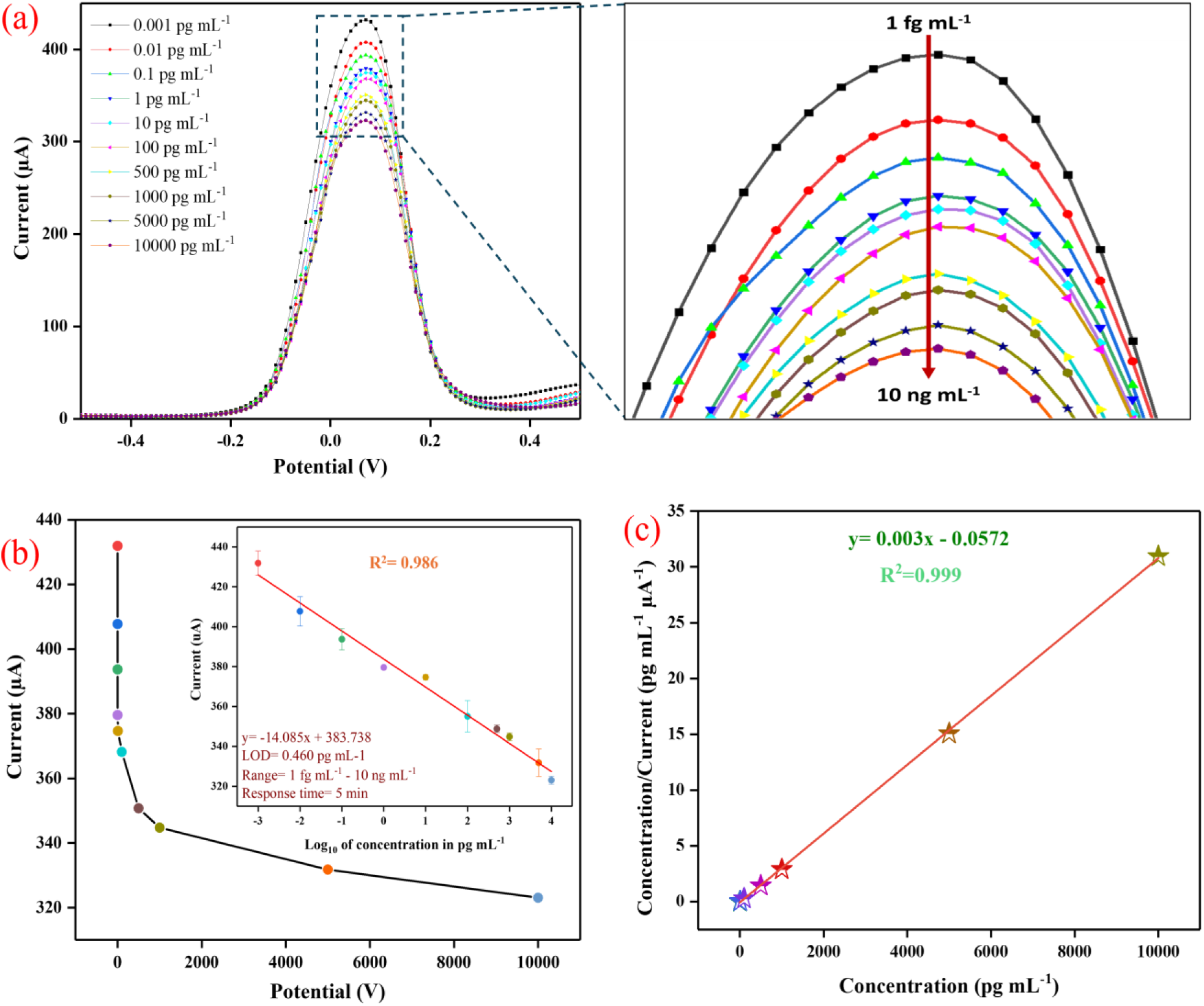
(a) Illustration of the electrochemical response of the sensing platform towards IL-6 concentrations using DPV (the enlarged view is displayed adjacent to it); (b) a calibration plot obtained using DPV techniques between the anodic peak current magnitude and the log_10_ of the IL-6 concentration; and (c) Hanes-Wolf plot.

As depicted in **Fig. 9(b)**, under optimized assay conditions, the change in current (ΔI) exhibited a highly linear relationship with IL-6 concentration across the range of 1 fg/mL to 10 ng/mL, with a linear regression coefficient (R^2^) of 0.986, confirming excellent analytical linearity. The sensitivity of the aptasensor was determined from the slope of the calibration curve to be 2.162 μA/[log_10_(ng/mL)] cm^-2^, reflecting the high responsiveness of the MCH/Apt-IL-6/g-C_3_N_4_@CS/Au-SPE platform toward IL-6. The computed limit of detection (LOD) was 0.460 pg mL⁻¹ (S/N = 3), which falls well within the clinically relevant range for IL-6 in human serum during early-stage sepsis and systemic inflammatory conditions, where IL-6 levels typically range from a few pg/mL to several hundred pg/mL. Beyond this range, the current stabilizes as higher concentrations are added. Eqn. (5) yields a linear regression coefficient (R^2^) of 0.986, affirming this relationship. The binding affinity of the aptasensor was evaluated by constructing a Hanes-Woolf plot between the IL-6 concentration and the corresponding current response of the MCH/Apt-IL-6/g-C_3_N_4_@CS/Au-SPE aptaelectrode [**Fig. 9(c)**]. The association constant (Ka) and dissociation constant (Kd) were derived from the plot, confirming the strong, specific binding affinity of the immobilized IL-6 aptamer for its target. A comparison of the analytical performance of the developed aptasensor with previously reported IL-6 biosensors is presented in **Table 3**, demonstrating that the MCH/Apt-IL-6/NC/Au-SPE aptasensor exhibits competitive or superior performance across linear detection range, sensitivity, and LOD.

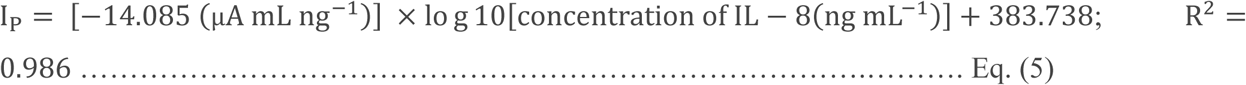

**Table 3:**
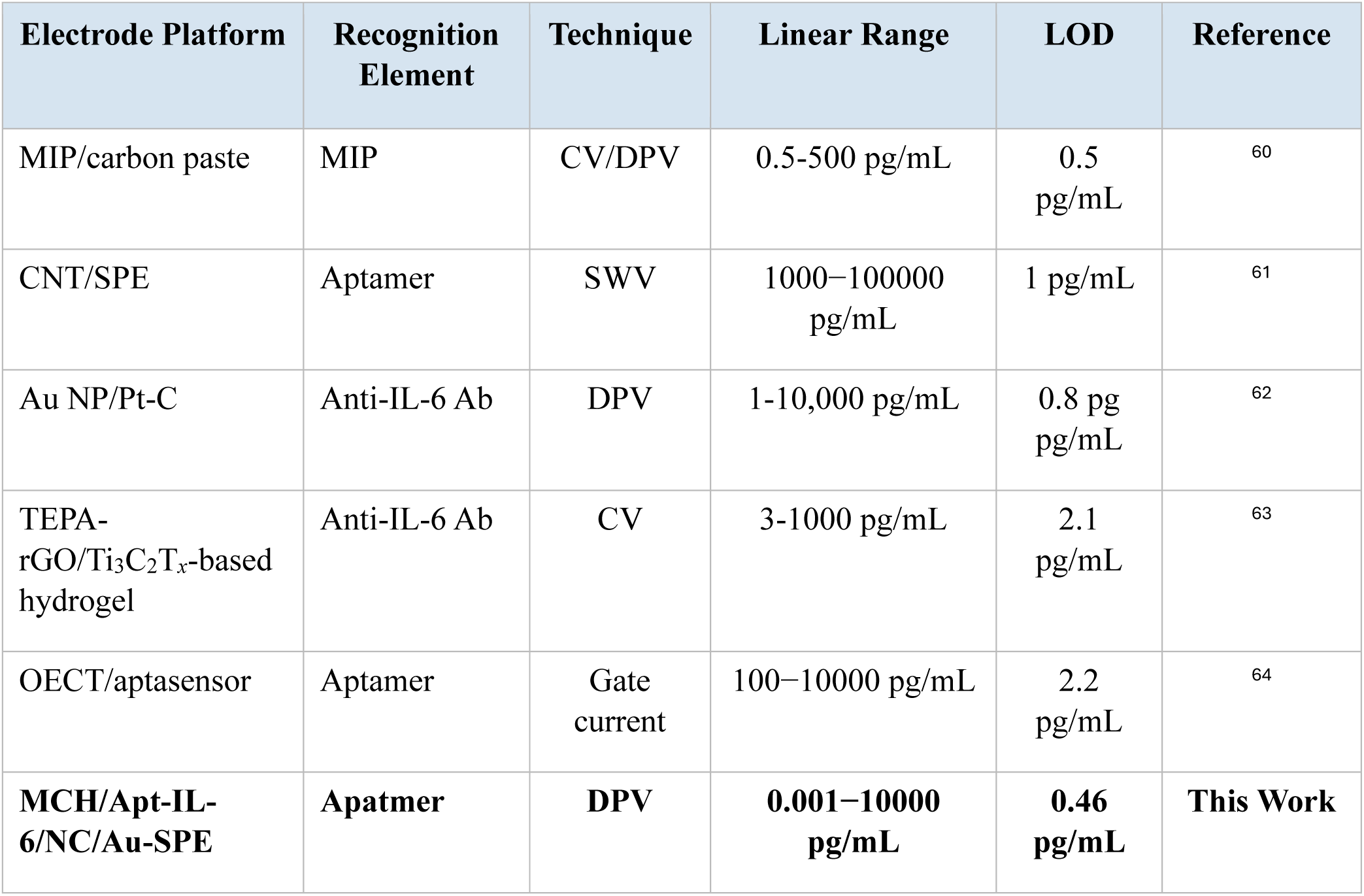
Previously reported IL-6 biosensors.

#### 3.4.6 Selectivity and Reproducibility Studies

Selectivity is among the most critical performance attributes of any electrochemical biosensor, given the complex composition of real biological matrices. The selectivity of the MCH/Apt-IL-6/g-C_3_N_4_@CS/Au-SPE aptasensor was evaluated by assessing its electrochemical response against a panel of common biological interferents present in human serum, including ascorbic acid, cysteine, glucose, glycine, and urea, each tested at a fixed concentration using DPV [**Fig. 10 (a&b)**]. The aptasensor exhibited significantly greater current suppression in the presence of IL-6 than with any of the tested interferents, which produced negligible changes in the DPV signal. These results confirm the outstanding selectivity of the aptasensor toward IL-6, which is primarily governed by the high structural specificity of the IL-6 aptamer for its cognate target, ensuring minimal cross-reactivity with structurally distinct biomolecules in complex sample matrices.

**Figure 10:**
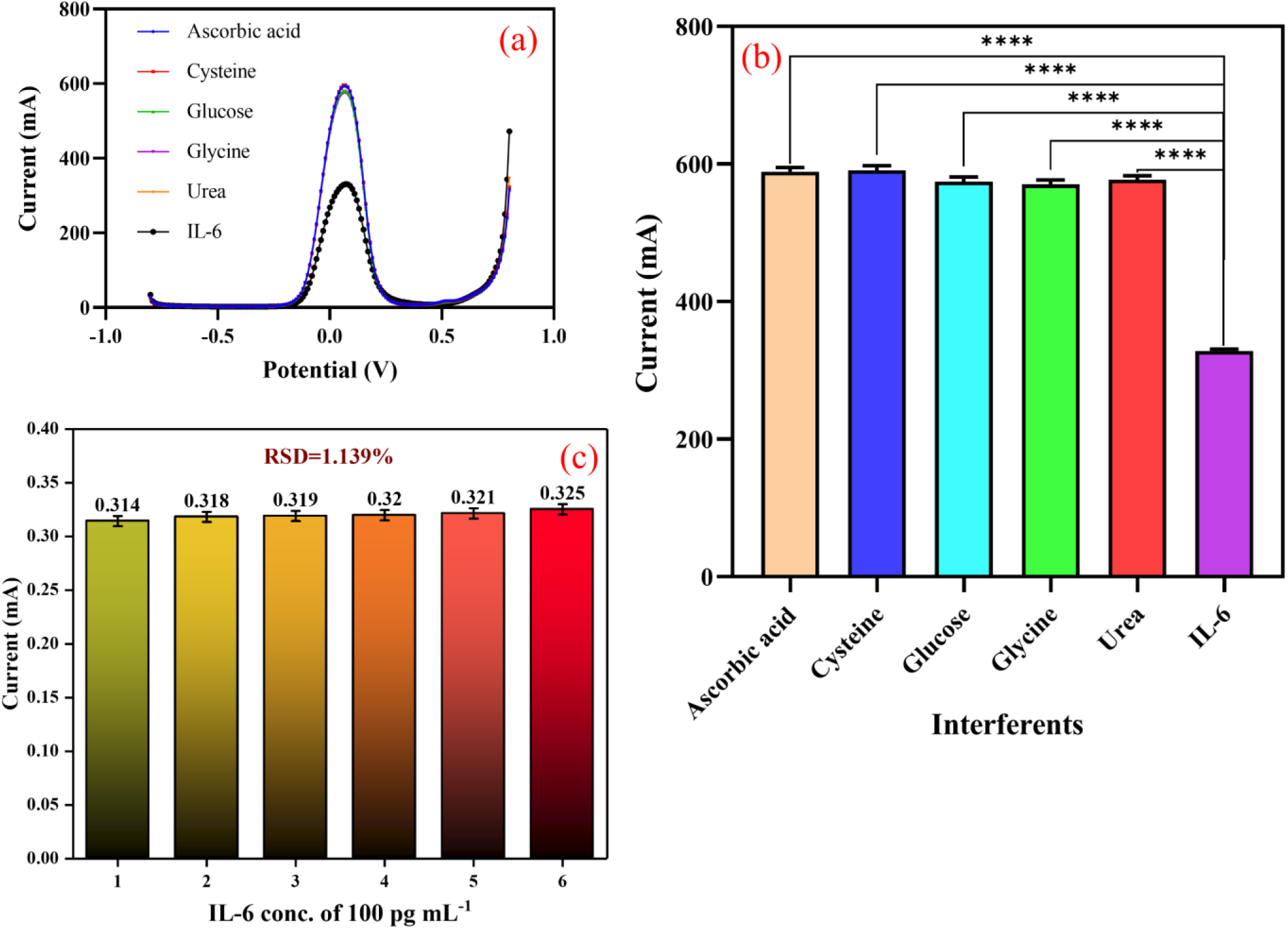
**(a)** and **(b)** Interferent study of different fabricated electrodes, and **(c)** repeatability study.

Reproducibility was assessed by fabricating six independent MCH/Apt-IL-6/g-C_3_N_4_@CS/Au-SPE aptaelectrodes under identical experimental conditions and recording their DPV responses individually [**Fig. 10(c)**]. The aptasensor demonstrated excellent inter-electrode reproducibility with an RSD of 1.139%, confirming the consistency and reliability of the fabrication protocol. These low RSD values validate the robustness of the g-C_3_N_4_@CS hydrogel modification approach and the aptamer immobilization strategy for producing reproducible aptasensor platforms.

#### 3.4.7 Real Sample Analysis

Analyzing IL-6 protein in biological samples is becoming increasingly necessary. The practical applicability of the MCH/Apt-IL-6/g-C_3_N_4_@CS/Au-SPE aptasensor was validated by detecting IL-6 in spiked artificial human serum, a physiologically representative complex matrix. The electrochemical response of the aptaelectrode toward spiked serum samples was recorded by DPV using a redox couple. A signal-off response was consistently observed, where the DPV peak current decreased linearly with IL-6 concentration. This phenomenon was replicated in standard PBS calibration experiments. The recovery percentage values from spiked serum experiments are presented in **Table 4**. Based on the results, it is suggested that the MCH/Apt-IL-6/g-C_3_N_4_@CS/Au-SPE aptasensor can quantitatively assess IL-6 in clinically relevant sample matrices, indicating its applicability for the detection of sepsis and inflammatory biomarkers in a point-of-care context.

**Table 4:** IL-6 recovery rates determined using the MCH/Apt-IL-6/g-C_3_N_4_@CS/Au-SPE aptaelectrode with real spiked samples.

| S. No | IL-6 conc. (pg mL<sup>-1</sup>) | Peak current (mA) in standard samples | Peak current (mA) in standard samples | % Recovery |
| --- | --- | --- | --- | --- |
| 1 | 0.001 | 431.896 | 415.78 | 96.26855 |
| 2 | 0.01 | 407.723 | 369.14 | 90.53696 |
| 3 | 0.1 | 393.693 | 328.188 | 83.3614 |
| 4 | 1 | 379.568 | 325.439 | 85.73931 |
| 5 | 10 | 374.638 | 324.396 | 86.58919 |
| 6 | 100 | 355.015 | 323.258 | 91.05474 |
| 7 | 500 | 348.759 | 321.173 | 92.09024 |
| 8 | 1000 | 344.777 | 317.855 | 92.19147 |
| 9 | 5000 | 331.79 | 316.717 | 95.45707 |
| 10 | 10000 | 323.069 | 307.332 | 95.1289 |

## 4. Conclusion

In conclusion, a biofunctional, label-free electrochemical aptasensor based on a g-C3N4-incorporated CS hydrogel-modified gold screen-printed electrode (MCH/Apt-IL-6/g-C_3_N_4_@CS/Au-SPE) was successfully developed for the ultrasensitive detection of IL-6, a clinically important inflammatory biomarker closely associated with sepsis progression and SCM. The engineered g-C_3_N_4_@CS heterointerface was formed via electrostatic interactions between negatively charged g-C3N4 nanosheets and protonated amino groups of CS, yielding a porous, electrochemically active, and biocompatible hydrogel network that facilitated efficient electron transfer and enhanced aptamer immobilization. Comprehensive physicochemical, electrochemical, and biological investigations confirmed the successful fabrication and functionality of the sensing platform. Biocompatibility evaluation using MTT assay and confocal fluorescence imaging demonstrated that the hydrogel exhibited excellent cytocompatibility at lower concentrations, particularly at 5 mg/mL, where cells maintained normal morphology, intact cytoskeletal organization, preserved nuclear integrity, and active mitochondrial distribution comparable to untreated controls. In contrast, higher concentrations (10 mg/mL) induced noticeable cellular stress and reduced viability, establishing 5 mg/mL as the optimal concentration for biosensor fabrication and biomedical applicability. Under optimized experimental conditions, the developed aptasensor exhibited remarkable analytical performance, including an ultrawide linear detection range from 1 fg/mL to 10 ng/mL, high sensitivity of 2.162 μA/[log_10_(ng/mL)] cm^-2^, a low detection limit of 0.460 pg/mL, and excellent linearity (R^2^ = 0.986). Furthermore, the sensor displayed high selectivity toward common biological interferents, including ascorbic acid, cysteine, glucose, glycine, and urea, along with outstanding reproducibility (RSD = 1.139%). Validation in spiked human serum samples further demonstrated the practical reliability and robustness of the proposed sensing strategy for clinical biofluid analysis. Compared with conventional immunoassays, the proposed aptasensing platform offers several significant advantages, including rapid, label-free detection; antibody-free molecular recognition; simple, scalable fabrication; disposable screen-printed electrode integration; and cost-effective operation. Collectively, these findings highlight the strong potential of the developed g-C_3_N_4_-hydrogel heterointerface as a next-generation electrochemical biosensing platform for rapid point-of-care diagnostics, early screening for SCM, and real-time monitoring of inflammatory biomarkers in emergency healthcare settings. Future studies integrating artificial intelligence-assisted signal analysis, wearable sensing modules, and IoT-enabled healthcare systems may further accelerate its translation into intelligent, clinically deployable diagnostic technologies.

## Data availability

The data supporting this article are included in the SI.

## Credit authorship contribution statement

**Pranjal Agarwaal:** Writing**-** original draft, Software, Methodology, Investigation**. Amit K. Yadav:** Conceptualization, Project administration, Software, Methodology, Supervision, Writing - original draft, Writing - review & editing. **Ankur Singh:** Visualization, Methodology, Investigation. **Sumit K. Yadav**-Software, Data curation, Writing- original draft. **N V S Praneeth**- Writing - review & editing. **Dhiraj Bhatia:** Writing - review & editing, Supervision, Project administration, Funding acquisition, Conceptualization.

## Acknowledgements

The authors acknowledge the infrastructure and support provided by the Indian Institute of Technology Gandhinagar, including facilities at the Central Instrumentation Facility (CIF). PA thanks the AICTE, Ministry of Education, Govt. of India, for the GATE fellowship. AKY thanks the Anusandhan National Research Foundation (ANRF), Govt. of India, for the National Post Doctoral Fellowship (N-PDF). DB acknowledges financial support from the Ministry of Education, Government of India. The authors also thank the library services at IIT Gandhinagar for providing access to scientific literature and plagiarism detection tools. We acknowledge the use of BioRender for preparing the illustrations.

---

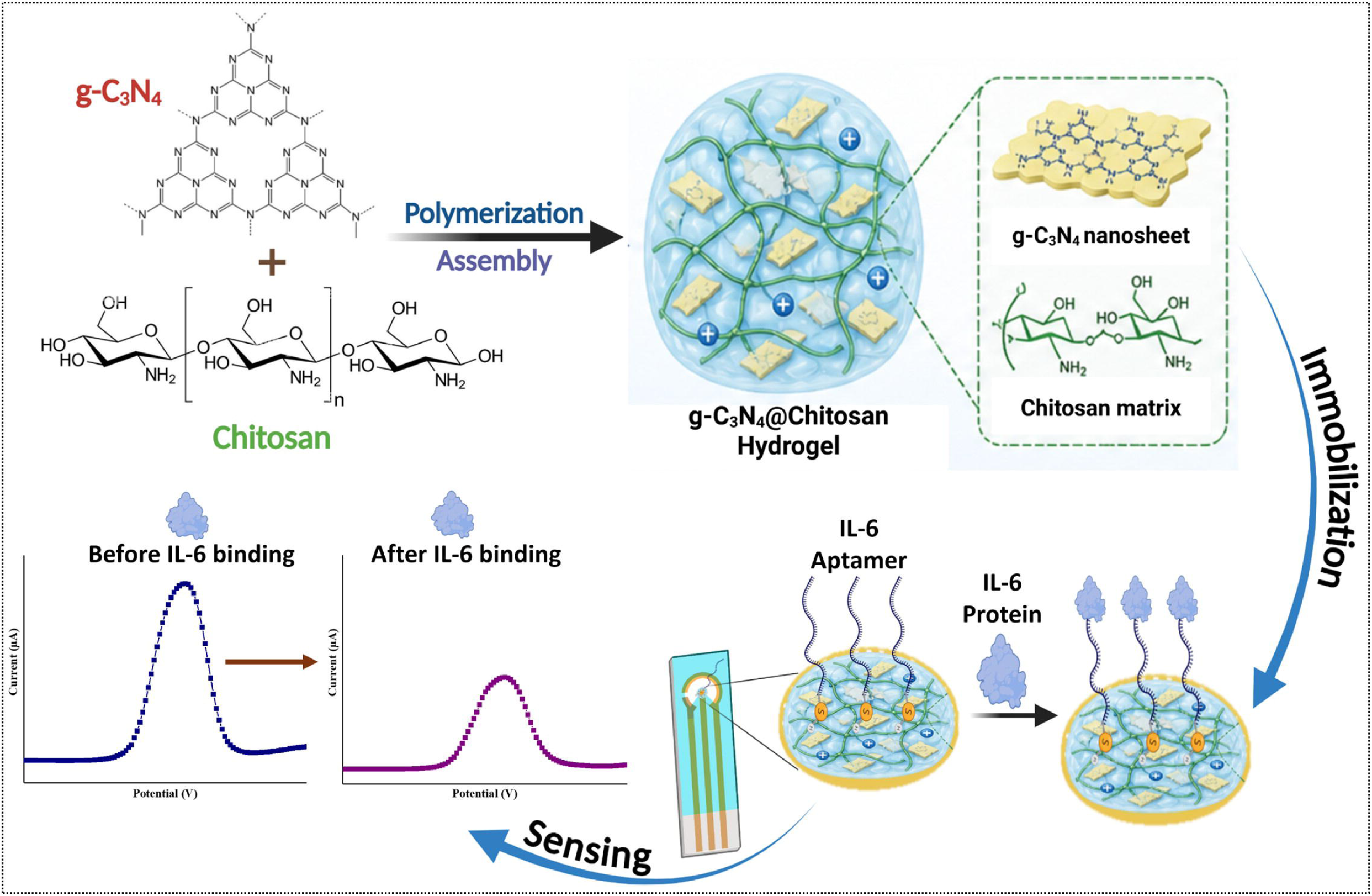

