## Supporting Data for "Biofunctional 2D Graphitic Carbon Nitride-Hydrogel Heterointerfaces for Electrochemical Detection of Interleukin-6 toward Septic Cardiomyopathy Diagnostics in Clinical Biofluids"

**[$]Authors Contributed Equally to this work**

***Corresponding Authors:**

### SUPPLEMENTARY INFORMATION

**Table S1**

| | |
|---|---|
| IL-6 aptamer sequence | [ThiC6]GGTGGCAGGAGGACTATTTATTTGCTTTTCT |

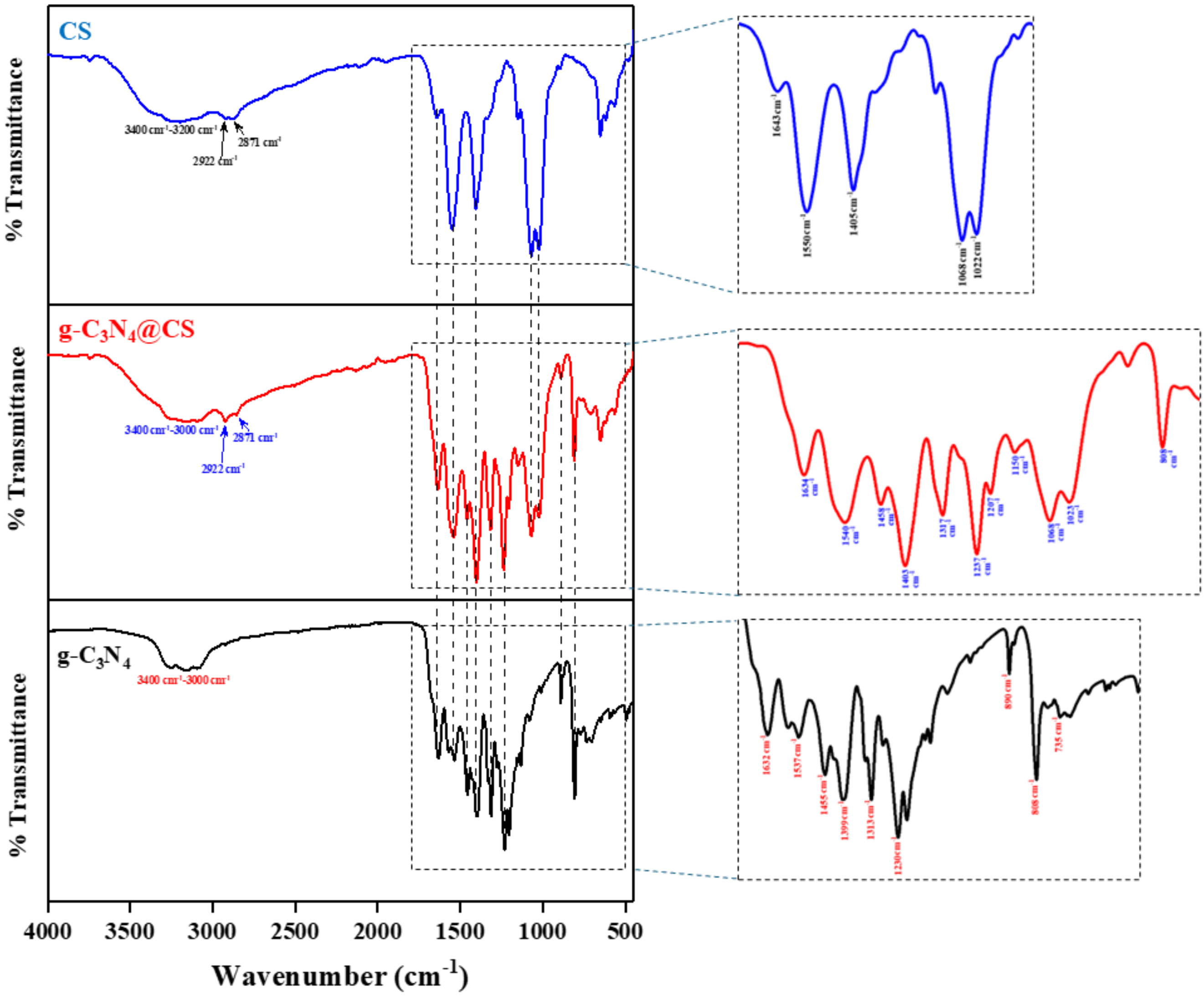


**Figure S1:** FT-IR spectra of g-$C_3N_4$, chitosan, and g-$C_3N_4$@CS hydrogel.

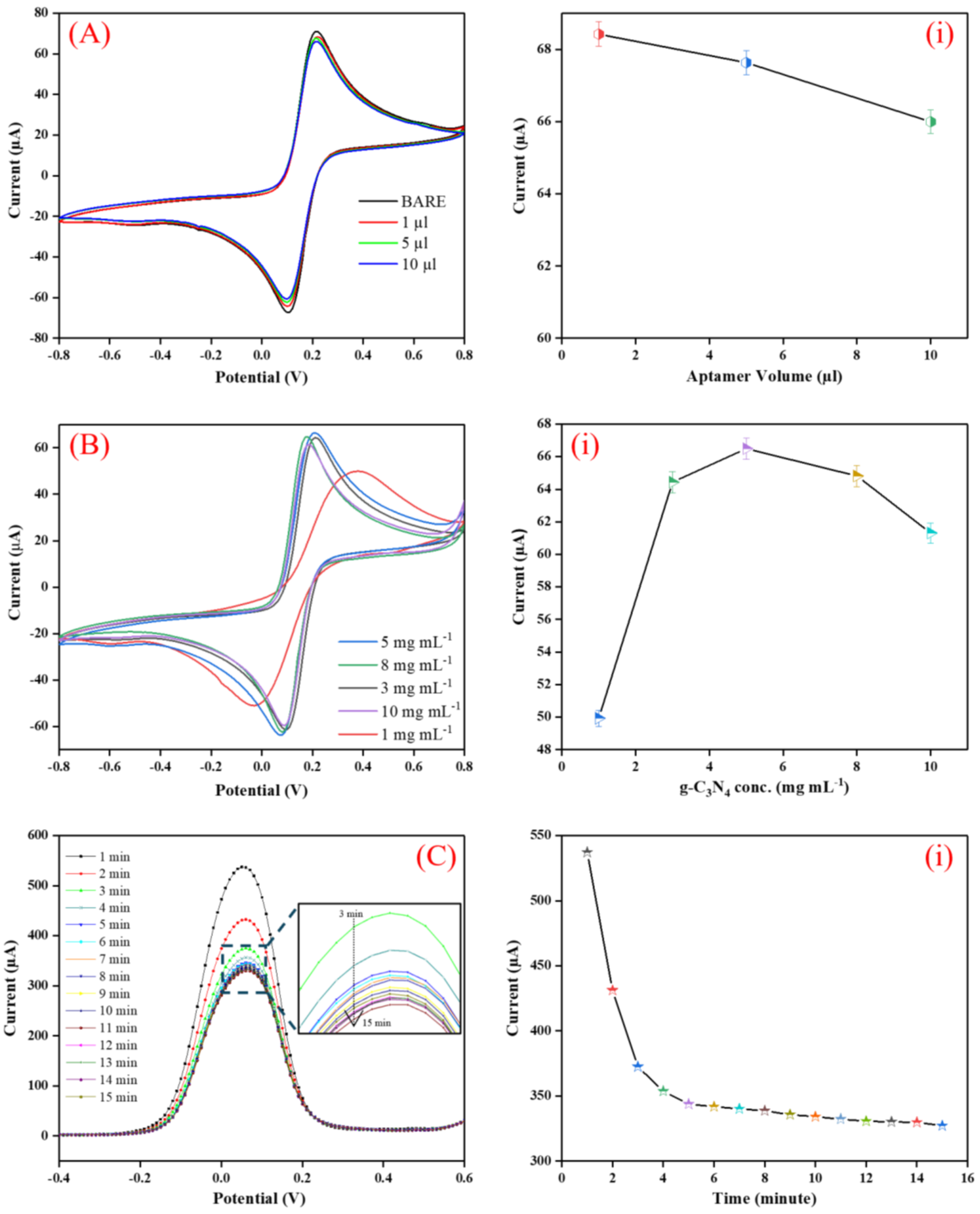


**Figure S2:** Optimized parameters for electrochemical sensing. [(A) & (i)] Aptamer volume; [(B) & (i)] g-$C_3N_4$ conc.; and [(C) & (i)] Response time

**Table S2:** Optimized parameters for IL-6 sensing

| Parameter | Range | Optimized value |
|---|---|---|
| **Aptamer volume** | 1-10 µL | 5 µL |
| **$g\text{-}C_3N_4$ concentration** | 1-10 mg/mL | 5 mg/mL |
| **Incubation time** | 1-15 min | 6 min |
| **$g\text{-}C_3N_4$ volume** | 1-5 µL | 5 µL |
| **MCH volume** | 1-5 µL | 5 µL |

**Kinetic Interface Studies: Quantitative Parameter Calculations**

To comprehensively characterize the electrochemical interface at each key stage of aptasensor fabrication, four critical kinetic parameters were calculated from the scan rate data: the diffusion coefficient (D), the electroactive surface area (Ae), the surface concentration of electroactive ionic species (γ*), and the heterogeneous electron transfer rate constant (Ks). These calculations were performed for both the g-$C_3N_4$@CS/Au-SPE electrode (before aptamer immobilization) and the fully assembled MCH/Apt-IL-6/g-$C_3N_4$@CS/Au-SPE aptaelectrode. The experimental conditions used throughout were: Au-SPE working electrode (geometric area A = 0.25 $cm^2$), PBS (pH 7.5) containing 5 mM $[Fe(CN)_6]^{3-/4-}$ (C = $5 \times 10^{-6}$ mol/$cm^3$), scan rate v = 50 mV/s (0.05 V/s), T = 298 K, n = 1 (one electron transfer per $Fe^{2+}/Fe^{3+}$ event), F = 96,485 C/mol, R = 8.314 J $mol^{-1}$ $K^{-1}$.

The peak currents measured at 50 mV/s from the scan rate CV experiments were: for the g-$C_3N_4$@CS/Au-SPE, Ipa = 87.7 µA and Ipc = 90.1 µA; for the MCH/Apt-IL-6/g-$C_3N_4$@CS/Au-SPE, Ipa = 43.7 µA and Ipc = 40.7 µA.

**(i) Diffusion Coefficient (D) - Randles-Ševčík Equation**

The diffusion coefficient (D) of $[Fe(CN)_6]^{3-/4-}$ at each electrode interface was determined using the Randles-Ševčík equation, which relates the peak current (Ip) to the square root of the scan rate through the diffusion coefficient, electrode area, and analyte concentration:

$$Ip = (2.69 \times 10^5) \times n^{3/2} \times D^{1/2} \times v^{1/2} \times A \times C \qquad \text{............... (Eq. i)}$$

Rearranging for D:

$$D = [\, Ip \,/\, (2.69 \times 10^5 \times n^{3/2} \times v^{1/2} \times A \times C) \,]^2 \qquad \text{............... (Eq. ii)}$$

Substituting the measured values for the g-$C_3N_4$@CS/Au-SPE (Ipa = $87.7 \times 10^{-6}$ A, n = 1, ν = 0.05 V/s, A = 0.25 $cm^2$, C = $5 \times 10^{-6}$ mol/$cm^3$):

D (g-$C_3N_4$@CS/Au-SPE) = $[87.7 \times 10^{-6} / (2.69 \times 10^5 \times 1 \times 0.2236 \times 0.25 \times 5 \times 10^{-6})]^2$ = $1.361 \times 10^{-6}$ $cm^2$/s = $136.10 \times 10^{-8}$ $cm^2$/s

Substituting the measured values for the MCH/Apt-IL-6/g-$C_3N_4$@CS/Au-SPE (Ipa = $48.0 \times 10^{-6}$ A):

D (MCH/Apt-IL-6/g-$C_3N_4$@CS/Au-SPE) = $[43.7 \times 10^{-6} / (2.69 \times 10^5 \times 1 \times 0.2236 \times 0.25 \times 5 \times 10^{-6})]^2$ = $3.38 \times 10^{-7}$ $cm^2$/s = $33.80 \times 10^{-8}$ $cm^2$/s

The lower D value for the fully assembled MCH/Apt-IL-6/g-$C_3N_4$@CS/Au-SPE ($33.80 \times 10^{-8}$ $cm^2$/s) compared to the g-$C_3N_4$@CS/Au-SPE ($136.10 \times 10^{-8}$ $cm^2$/s) clearly reflects the steric and electrostatic barriers introduced by the aptamer and MCH layers, which restrict the diffusion of $[Fe(CN)_6]^{3-/4-}$ toward the electrode surface. The aptamer strand, being a large, negatively charged nucleic acid molecule, electrostatically repels the anionic redox probe, while the hydrophobic MCH alkyl chain acts as a physical diffusion barrier at bare gold sites, together creating a dual hindrance mechanism that is quantitatively captured by the reduction in D.

**(ii) Electroactive Surface Area (Ae)**

The effective electroactive surface area (Ae) available for redox probe interaction was calculated from the Randles-Ševčík equation using the published literature diffusion coefficient of $[Fe(CN)_6]^{3-/4-}$ in PBS as the reference value, enabling a consistent cross-electrode comparison:

$$Ae = Ip / (2.69 \times 10^5 \times n^{3/2} \times D\_lit^{1/2} \times v^{1/2} \times C) \quad \text{…………… (Eq. iii)}$$

For the g-$C_3N_4$@CS/Au-SPE:

Ae (g-$C_3N_4$@CS/Au-SPE) = $87.7 \times 10^{-6} / (2.69 \times 10^5 \times 1 \times 5 \times 10^{-6} \times 2.588 \times 10^{-3} \times 0.2236)$ = 0.1126 $cm^2$

For the MCH/Apt-IL-6/g-$C_3N_4$@CS/Au-SPE:

Ae (MCH/Apt-IL-6/g-$C_3N_4$@CS/Au-SPE) = $43.7 \times 10^{-6} / (2.69 \times 10^5 \times 1 \times 5 \times 10^{-6} \times 2.588 \times 10^{-3} \times 0.2236)$ = 0.0561 $cm^2$

The g-$C_3N_4$@CS/Au-SPE exhibited a larger electroactive surface area (0.1126 $cm^2$) compared to the fully assembled aptaelectrode (0.0561 $cm^2$). This reduction in Ae upon aptamer immobilization and MCH passivation is expected and desired: the aptamer strands and MCH molecules physically occupy and passivate a fraction of the electroactive gold and hydrogel surface, reducing the number of sites accessible to the $[Fe(CN)_6]^{3-/4-}$ redox probe. At the same time, the larger Ae of the g-$C_3N_4$@CS/Au-SPE relative to the geometric electrode area (0.25 $cm^2$ = 25 $mm^2$) confirms that the g-$C_3N_4$@CS porous hydrogel coating contributes significant internal surface area above the flat geometric surface, providing abundant sites for aptamer immobilization and explaining the improved biosensor sensitivity.

**(iii) Surface Concentration of Electroactive Species (γ*)**

The surface concentration of electroactive ionic species (γ*, mol $cm^{-2}$) adsorbed or accessible at the electrode interface was calculated using the Brown-Anson equation, which relates the peak current to the surface coverage of electroactive species under surface-confined or quasi-confined conditions:

$$Ip = (n^2 F^2 \gamma^* A v) / (4 R T) \quad \text{.............. (Eq. iv)}$$

Rearranging to solve for γ*:

$$\gamma^* = (4 R T \times Ip) / (n^2 F^2 A v) \quad \text{.............. (Eq. v)}$$

Substituting constants and measured values for the g-$C_3N_4$@CS/Au-SPE:

γ* (g-$C_3N_4$@CS/Au-SPE) = $(4 \times 8.314 \times 298 \times 87.7 \times 10^{-6}) / (1^2 \times 96485^2 \times 0.25 \times 0.05)$ = $7.47 \times 10^{-9}$ mol/$cm^2$ = $0.747 \times 10^{-8}$ mol/$cm^2$

For the MCH/Apt-IL-6/g-$C_3N_4$@CS/Au-SPE:

γ* (MCH/Apt-IL-6/g-$C_3N_4$@CS/Au-SPE) = $(4 \times 8.314 \times 298 \times 43.7 \times 10^{-6}) / (1^2 \times 96485^2 \times 0.25 \times 0.05)$ = $3.72 \times 10^{-9}$ mol/$cm^2$ = $0.372 \times 10^{-8}$ mol/$cm^2$

The higher γ* for the g-$C_3N_4$@CS/Au-SPE ($0.747 \times 10^{-8}$ mol/$cm^2$) compared to the MCH/Apt-IL-6/g-$C_3N_4$@CS/Au-SPE ($0.372 \times 10^{-8}$ mol/$cm^2$) indicates that more ionic redox species can access and interact with the hydrogel electrode surface before aptamer and MCH passivation. The nitrogen-rich surface of g-$C_3N_4$ and the protonated amine groups of chitosan create a positive surface environment that electrostatically attracts and locally concentrates the negatively charged $[Fe(CN)_6]^{3-/4-}$ species, increasing effective γ* at the g-$C_3N_4$@CS/Au-SPE interface. After aptamer immobilization and MCH blocking, partial neutralization of surface charge and steric exclusion reduce the local concentration of accessible electroactive species, as evidenced by the lower γ* of the assembled aptasensor.

**(iv) Heterogeneous Electron Transfer Rate Constant (Ks)**

The heterogeneous electron transfer rate constant (Ks, $s^{-1}$), a quantitative measure of the intrinsic speed of electron exchange between the electrode surface and the redox species in solution, was evaluated using the Laviron equation, which relates Ks to the peak-to-peak separation (ΔEp) observed in the cyclic voltammogram:

$$Ks = m \times n \times F \times v \ / \ (R \times T) \qquad \text{.............. (Eq. vi)}$$

where m represents the peak-to-peak separation of potentials (ΔEp, in V), n is the number of electrons transferred, F is the Faraday constant, ν is the scan rate (V/s), R is the universal gas constant, and T is the absolute temperature (K).

For the g-$C_3N_4$@CS/Au-SPE (ΔEp = 68 mV = 0.068 V at 50 mV/s):

Ks (g-$C_3N_4$@CS/Au-SPE) = $0.068 \times 1 \times 96485 \times 0.05 \ / \ (8.314 \times 298) = 0.132\ s^{-1}$

For the MCH/Apt-IL-6/g-$C_3N_4$@CS/Au-SPE (ΔEp = 143 mV = 0.143 V):

Ks (MCH/Apt-IL-6/g-$C_3N_4$@CS/Au-SPE) = $0.143 \times 1 \times 96485 \times 0.05 \ / \ (8.314 \times 298) = 0.278\ s^{-1}$

The g-$C_3N_4$@CS/Au-SPE electrode shows a smaller ΔEp (68 mV) compared to the assembled aptasensor (143 mV), which, within the Laviron framework, reflects faster, kinetically less impeded electron transfer at the hydrogel surface before biological blocking layers are applied. The larger ΔEp of the MCH/Apt-IL-6/g-$C_3N_4$@CS/Au-SPE corresponds to a greater kinetic departure from electrochemical reversibility, consistent with the steric and electrostatic barriers imposed by the aptamer and MCH layers. This progressive increase in ΔEp with each surface modification step from near-reversible (68 mV) to quasi-reversible (143 mV) provides a clear qualitative and quantitative indicator of each successive surface blocking event and confirms the integrity of the aptasensor assembly process.

**Calculation of the number of molecules in IL6 at LOD of 0.406 pg/mL**

Assuming the molecular weight (MW) of IL-6 is approximately 21 kDa = 21,000 g/mol.

Step 1: Convert concentration into grams

$$0.406 \text{ pg/mL} = 0.406 \times 10^{-12} \text{ g/mL}$$

Step 2: Calculate the number of moles

$$n = \frac{m}{M} = \frac{0.406 \times 10^{-12}}{21000}$$

$$n = 1.93 \times 10^{-17} \text{ mol/mL}$$

Step 3: Convert moles to the number of molecules

Using Avogadro's number:

$$N_A = 6.022 \times 10^{23} \text{ molecules/mol}$$

$$N = (1.93 \times 10^{-17})(6.022 \times 10^{23})$$

$$N = 1.16 \times 10^{7} \text{ molecules/mL}$$

Final Answer

At an LOD of 0.406 pg/mL, the sample contains approximately: $1.16 \times 10^{7}$ molecules/mL
